# Columnar localization and laminar origin of cortical surface electrical potentials

**DOI:** 10.1101/2021.08.16.456540

**Authors:** Vyassa L. Baratham, Maximilian E. Dougherty, Peter Ledochowitsch, Michel M. Maharbiz, Kristofer E. Bouchard

**Affiliations:** Biological Systems and Engineering Division; Lawrence Berkeley National Laboratory, Berkeley CA 94720; Dept. of Physics, UC Berkeley, Berkeley CA 94720; Allen Institute for Brain Science, Seattle, WA 98109; Center for Neural Engineering and Prosthesis, UCSF-UCB; Dept. of Electrical Engineering and Computer Science, UC Berkeley, Berkeley CA 94720; Helen Wills Neuroscience Institute and Redwood Center for Theoretical Neuroscience, UC Berkeley, Berkeley CA 94720; Computational Research & Biological Systems and Engineering Divisions, Lawrence Berkeley National Laboratory, Berkeley CA 94720

**Author notes:** These authors contributed equally to this work (Vyassa L. Baratham, Maximilian E. Dougherty).

**Keywords:** origins of ECoG, biophysical simulations, auditory cortex, cortical column

## Abstract

Electrocorticography (ECoG) methodologically bridges basic neuroscience and understanding of human brains in health and disease. However, the localization of ECoG signals across the surface of the brain and the spatial distribution of their generating neuronal sources are poorly understood. To address this gap, we recorded from rat auditory cortex using customized μECoG, and simulated cortical surface electrical potentials with a full-scale, biophysically detailed cortical column model. Experimentally, μECoG-derived auditory representations were tonotopically organized and signals were anisotropically localized to ≤±200 μm, i.e., a single cortical column. Biophysical simulations reproduce experimental findings, and indicate that neurons in cortical layers V and VI contribute ∼85% of evoked high-gamma signal recorded at the surface. Cell number and synchronicity were the primary biophysical properties determining laminar contributions to evoked μECoG signals, while distance was only a minimal factor. Thus, evoked μECoG signals primarily originate from neurons in the infragranular layers of a single cortical column.

**In Brief:** Baratham et al., investigated the localization and origins of sensory evoked ECoG responses. They experimentally found that ECoG responses were anisotropically localized ≤±200 μm, i.e., a single cortical column. Biophysically detailed simulations revealed that neurons in layers V &VI were the primary sources of evoked ECoG responses, in contrast to common thinking.

**Highlights:** Evoked μECoG signals are localized on the surface to a cortical column.

Neurons in cortical layers V and VI constitute the vast majority of the signal recorded at the surface.

Different laminar contributions to ECoG signal are driven by cell density and synchronicity.

## Introduction

The brain is composed of many neuronal microcircuits (e.g., cortical columns) that each perform specific computations, but are simultaneously integrated into larger networks^1,2^. Microscale measurements (e.g., whole cell and extracellular electrophysiology, Ca^2+^ imaging, etc.,) which investigate the activity of individual neurons and small neuronal populations have yielded insight into microcircuit mechanisms of local computations. At the same time, macroscale measurements (e.g., fMRI) have revealed principles of global processing in entire brain areas^3^. However, much less is known about how local neural processing is organized and coordinated across distributed brain networks. The relative paucity of knowledge about mammalian mesoscale (i.e., intermediate) cortical functioning can be attributed to the difficulty of simultaneously measuring the activity of 100s-1000s of functionally distinct sites over large spatial scales with sufficient resolution to resolve properties of local neuronal populations^4,5^. Measuring distributed cortical function is critical to understanding the computations giving rise to complex perceptions and behaviors^6–8^. High-density micro-electrocorticography (μECoG) arrays record neuronal activity directly from the cortical surface, can be fabricated on flexible materials with tight spacing of many thousands of channels, and are minimally invasive^9–11^. Furthermore, because ECoG and μECoG are used in humans, it is a critical methodological bridge between basic neuroscience findings and our understanding of the human brain in health and disease^9^. However, the utilization of (μ)ECoG for basic neuroscience is impeded by a lack of understanding of the spatial localization of the recorded signals across the surface and the specific neuronal sources generating those signals.

ECoG records cortical surface electrical potentials (CSEPs), which, like all electrical signals in the brain, reflect a weighted superposition of all electrical sources surrounding the electrode^3,12^. Because of the 1/f^α^ fall-off of power with frequency of brain signals^3,12–14^, many studies of ECoG and local field potentials (LFP) focus on lower frequencies (e.g., <60 Hz), though it is becoming more common to utilize activity in the ‘high-gamma’ band (Hγ: 65-170 Hz)^7,15^. The motivation for using Hγ stems from the proposal that higher frequencies reflect a more spatially localized signal^3^. However, experimental estimates of spatial spread of correlations range over more than an order of magnitude, from several hundred micrometers to a few millimeters. Reports on the frequency dependence of spatial spread are similarly inconsistent^16–18^. Determining the spatial localization of evoked ECoG signals is critical for both interpreting the data as well as guiding the design of future devices.

An additional critical issue is understanding how ECoG signals arise during active neuronal processing, as opposed to baseline. During sensory processing, neuronal populations are activated with spatial correlations that partly depend on functional organization across cortex and the biophysics of signal propagation through cortical tissue^19,20^. In primary sensory cortices (e.g., A1), the functional organization of response properties varies smoothly across adjacent columns^21^. A cortical column in rodents has a radius of 200-500 μm (∼350 μm) and spans ∼1800-2100 μm in depth, including all six cortical layers, which are composed of a variety of neuron types^22,23^. While different excitatory and inhibitory cell types within columnar microcircuits play different roles in sensory computation, in general neurons within a column share similar sensory tuning properties^24,25^. At the same time, electrical fields spread passively through cortical tissue, diffusing the signal^20^. Thus, both passive spread of the electric field through the tissue and functional organization of neuronal populations are potential mechanisms dictating localization of evoked ECoG signals across the surface. However, which of the two is the dominant mechanism is unknown.

Most fundamentally, we lack biophysical understanding of the precise sources that generate cortical surface electrical potentials (CSEPs). To understand how intracortical local field potentials (LFPs) arise from transmembrane currents, several recent studies have utilized biophysically detailed models. In models of passive neurons (i.e., no action potential generation), the spatial spread of population LFP is determined by single-neuron morphology, the temporal correlation between sources (synchronicity), and the number (density) of sources^20,26^. Additionally, the distance of the sources to the electrode is critical, as the magnitude of a single source decays with distance. Reimann et al.^27^, showed that, in contrast to commonly held belief, intra-cortical LFPs contain a significant contribution from active membrane current (e.g., voltage gated Na^+^ currents generating action potentials). However, all these studies used only three neuron types: two pyramidal neurons (layer III or IV, plus layer V) and layer IV stellate cells. Unlike for intracortical LFPs, biophysically detailed simulations have not been used to understand CSEPs. For example, Miller et al.^14^, created a phenomenological model of the shape of power vs. frequency recorded by macro-electrode ECoG in humans, and suggested that synaptic inputs were the major contributors to broad-band CSEPs. However, that study did not focus on evoked activity and the qualitative model cannot reveal the anatomical origin of ECoG, nor the biophysical principles for that origin. The layers of a cortical column are composed of different numbers of neurons with distinct morphologies and synchronicity, and are by definition at different distances from the surface electrode^28^. Thus, a full-scale, biophysically detailed cortical column model that reproduces the frequency content of evoked cortical surface electrical potentials is required to determine the precise laminar and cellular origins of evoked CSEPs.

We hypothesized that evoked high-frequency (e.g., high-gamma) cortical surface electrical potentials are spatially localized to a cortical column and are primarily generated by action potentials from neurons in layers V/VI. To test this hypothesis, we combined direct experimentation with biophysical modeling. We fabricated custom designed μECoG devices with small, low-impedance electrodes^10^, and recorded evoked CSEPs over the surface of rat auditory cortex. We provide definitive evidence that evoked μECoG signals are tightly and anisotropically localized to ≤±200 μm across the cortical surface. Thus, μECoG is localized to a cortical column, and the localization is primarily set by the spatial scales of functional organization of the underlying cortical tissue. To gain insight into the laminar contributions to evoked cortical surface signals, we created a full-scale, biophysically detailed model of a cortical column^28^ which was able to qualitatively recreate the evoked spectrum observed in the experimental data. Further, this model predicted that both the frequency content and amplitude of CSEP responses should increase with increasing input amplitude, which was confirmed by novel experimental findings. Finally, interrogation of the model shows that neurons in cortical layers V and VI contribute the vast majority (∼85%) of the signal recorded at the surface because of their increased cell number and synchronicity. These results provide a new biophysical understanding of the spatial localization of cortical surface electrical potentials and indicate that evoked ECoG signals originate from neurons in infragranular layers.

## Results

We designed and fabricated lithographically-defined, high-resolution μECoG arrays with customizable electrode geometry and spacing^10,11,29^. Each contact on the μECoG arrays had a diameter of 40 μm (approximately the size of a cortical mini-column), and an impedance of 30 ± 10 kΩ (measured in 1x phosphate-buffered saline at 1 kHz)^10^. To ensure that we fully captured the range of functional time-scales associated with the diversity of neurobiological signals, we recorded wide band (2-12,000 Hz) electrophysiological data.

### Stimulus-evoked cortical surface electrical potentials exhibit large peaks in the high-gamma range

An example of a 128-channel μECoG grid is shown on the cortical surface (subdural placement) of an anesthetized rat in the photomicrograph in **Figure 1a**. In this preparation, we recorded large amplitude, fast cortical surface electrical potentials (CSEPs) in response to the presentation of auditory tone pips (Methods). We played back stimuli consisting of short (50 ms) pure tone pips with varying frequency and intensity (amplitude) at 5 recording locations in 4 rats. An example recording from a single electrode during four consecutive stimulus presentations is shown in **Figure 1b**. Here, the top panel depicts stimulus amplitude and frequency, the middle panel displays the (normalized) neural spectrogram of the full response, and the bottom panel shows the amplitude of the high-gamma component of the neural response. For each recording electrode and CSEP frequency component, we normalized (z-scored, relative to baseline statistics taken during the inter-stimulus interval) the time-varying amplitude for each CSEP frequency separately, removed untuned electrodes (those which exhibit no stimulus frequency selectivity), and computed the “best frequency” for each tuned electrode (Methods). The best frequency is the stimulus frequency which maximally drives activity on that electrode. **Figure 1c**-**d** show the average evoked potential **(Fig. 1c)** and neural spectrogram **(Fig. 1d)** derived from the recorded electrical potential in response to the electrode’s best frequency at a single amplitude (N = 25 trials). In both single trials **(Fig. 1b)**, as well as the average **(Figs. 1d)**, the electrode’s best frequency evoked large-amplitude, rapid CSEP deflections. The high-gamma (Hγ: 65-170 Hz) component of the evoked response was observed to be the most robust, often exceeding five standard deviations of the baseline in response to the best-frequency (e.g., second stimulation, **Fig. 1b**, BF).

**Figure 1.**
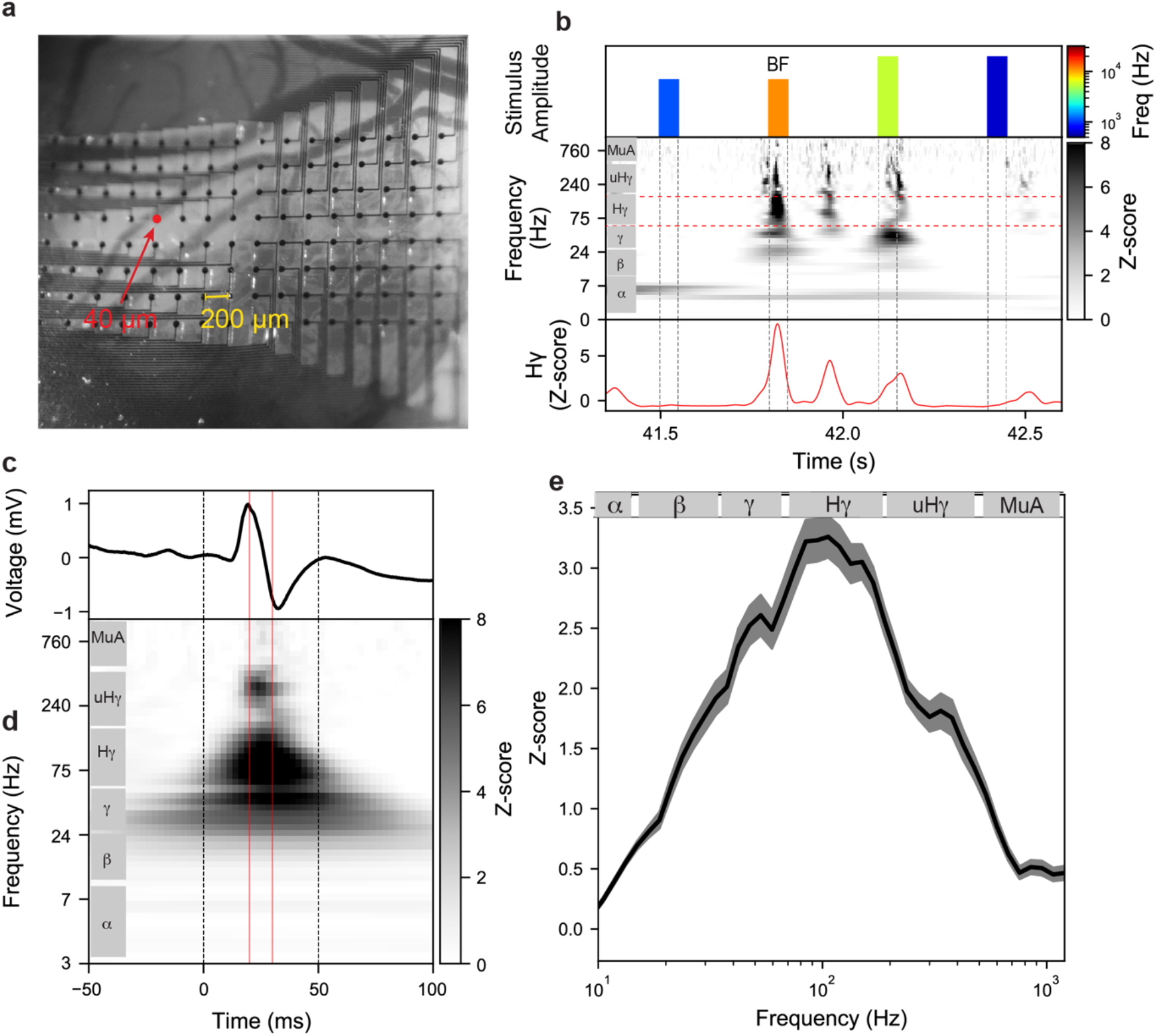
Stimulus-evoked cortical surface electrical potentials exhibit large peaks in the high-gamma range. **a**. Photomicrograph of an 8×16 μECoG grid (pitch: 200 μm, contact diameter: 40 μm) on the surface of rat primary auditory cortex (A1). **b**. Top: tone stimulus played during experimental recordings. Middle: z-scored spectral decomposition of single-trial evoked cortical surface electrical potentials from a single electrode. Bottom: High-gamma component of single-trial evoked cortical surface electrical potentials indicated by horizontal dashed lines in the middle panel. **c**. Trial-averaged evoked cortical surface electrical potential on one μECoG electrode in response to presentations of that electrode’s best tuned frequency. **d**. Trial-averaged neural spectrogram for the electrode shown in **c** in response to presentations of its best tuned frequency. Dashed vertical lines in **c** & **d** represent stimulus onset and offset. Red vertical lines in **c** & **d** correspond to the time window of extracted evoked response used for subsequent analysis. **e**. Grand-average (mean ± s.e.) z-scored amplitude as a function of frequency across all tuned electrodes (N = 333).

To summarize the frequency content of evoked CSEPs, we averaged across presentations of the best stimulus at one amplitude. For each electrode, we extracted mean z-scored responses across all neural frequency components in a ±5 ms window around the time of the peak high-gamma response (red vertical lines in **Fig. 1c-d**). We included all electrodes with a tuned response in the high-gamma band (N = 333 electrodes from 5 μECoG placements on auditory cortex in 4 rats). **Figure 1e** plots the averaged (N = 333 electrodes, mean ± s.e.) z-scored response as a function of frequency. On average, we found that evoked responses were unimodally peaked around the Hγ-band, with notable responses in the multi-unit activity range (MuA, >500 Hz).

### Robust frequency tuning and high-resolution tonotopic maps derived from μECoG

We next determined auditory receptive field properties and the spatial organization of evoked CSEPs. The plots of **Figure 2a-b** present frequency-response area (FRA) heat-maps derived from Hγ activity **(Fig. 2a)** and multi-unit activity (tMuA, after application of a threshold to the MuA band for event detection, Methods) **(Fig. 2b)** in response to these stimuli. In each FRA, pixels correspond to stimulus frequency-intensity pairings, and are colored according to the mean evoked z-scored signal (N = 25 stimuli for each), with blank spaces indicating un-tuned responses (see Methods). Data are displayed for several electrodes spanning 1.8mm anterior-posterior (AP) and 0.4 mm dorsal-ventral (DV) over rat primary auditory cortex (A1). We found that the response profiles exhibited clear frequency tuning and the canonical ‘V-shaped’ profiles expected of auditory cortical neurons^30–32^. Response profiles were largely similar between Hγ and tMuA. Additionally, there was a smooth gradation of best frequencies across the AP axis, with low-frequencies posterior, high-frequencies anterior, and similar best frequencies along the DV axis, suggestive of tonotopic organization.

**Figure 2.**
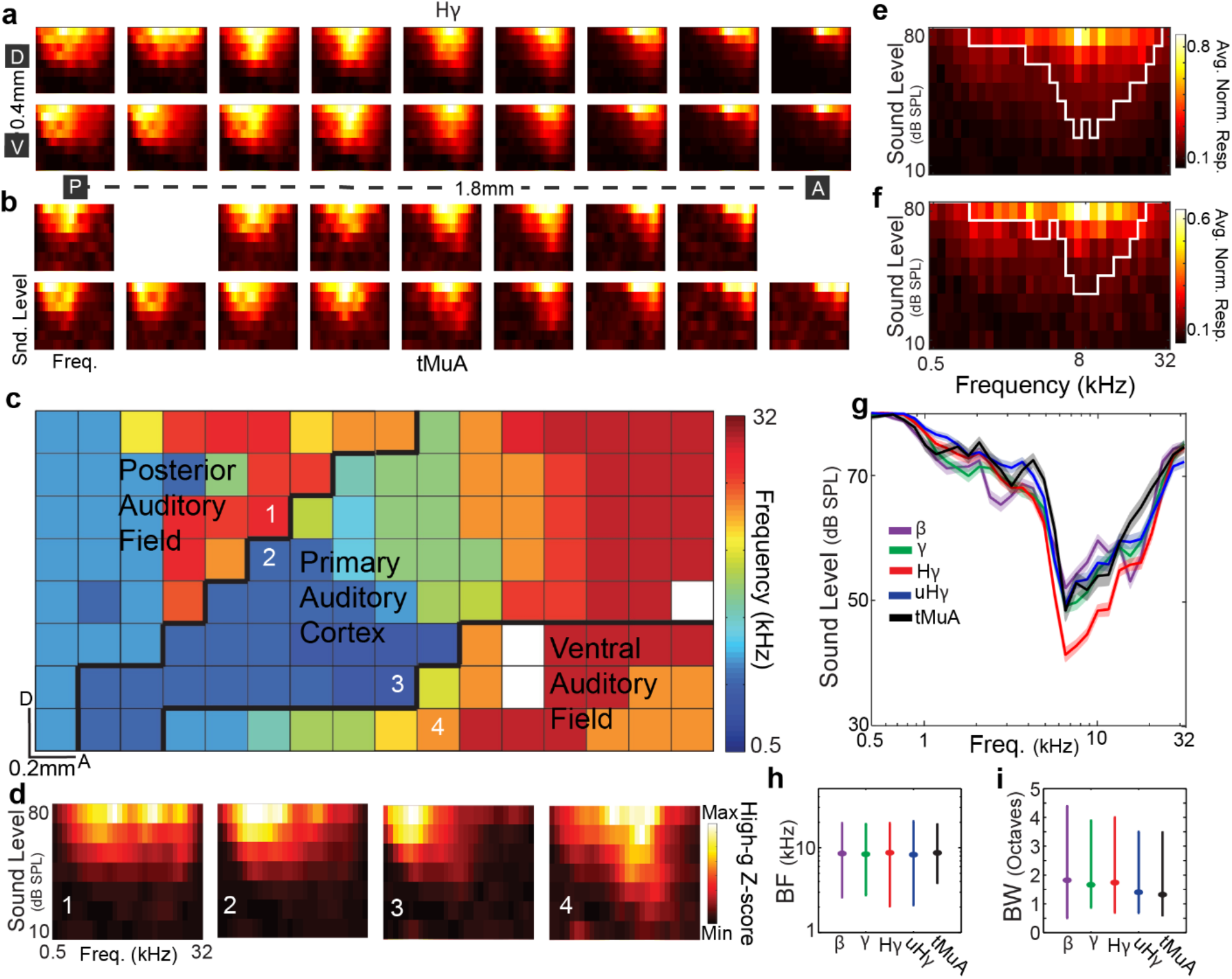
Robust frequency tuning and high-resolution tonotopic maps from μECoG. **a-b**. Frequency-response area (FRA) surfaces recorded from a μECoG array. Subplots correspond to responses of a single electrode and are organized according to electrode position on the grid/brain. In each subplot, pixels correspond to a stimulus frequency-intensity pairing, and are colored according to the mean evoked z-score. **a**. Hγ, **b**. tMuA. **c**. High-resolution tonotopic organization of multiple auditory cortical fields derived from Hγ activity. Each pixel is color coded according to that electrode’s best frequency. The 8×16 μECoG array displayed here covered multiple auditory cortical fields (A1, PAF, and VAF) and the approximate boundaries are demarcated (black lines). **d**. Differential tuning at neighboring electrodes. FRAs are plotted for four electrodes (numbered as in **c**) and show that neighboring electrodes (1 vs. 2; 3 vs. 4) can have different response properties. **e-f**. Average normalized response surface for all electrodes with significantly tuned Hγ (N = 333) and tMuA (N = 113) auditory responses. White line in each plot demarcates the FRA response boundaries. **g**. Across all tuned electrodes, the average (mean ± s.e.) FRA response boundaries for CSEP components (demarcated by colors) where similar. **h-i**. Distributions (25^th^-50^th^-75^th^ percentiles) of best-frequencies (**h**) and bandwidths (**i**) for all tuned responses for CSEP components.

As Hγ activity had the largest number of tuned channels, we first visualized tonotopic organization by coloring each electrode (pixel) according to its best frequency extracted from the Hγ-band **(Fig. 2c)**. The 8×16 μECoG array displayed here covered multiple auditory cortical fields [primary auditory cortex(A1), posterior auditory field (PAF), and ventral auditory field (VAF)] and the functionally defined boundaries are demarcated (black lines)^21,32^. Within a given auditory cortical field, there was a smooth gradation of best frequencies, with low-frequencies posterior and high-frequencies anterior. Tuning across the dorsal-ventral direction was largely similar within an auditory field. Interestingly, while frequency tuning generally varied smoothly as a function of distance between electrodes within an auditory area, we observed examples of different tuning at neighboring electrodes located in different auditory fields. For example, the FRAs for the electrodes demarcated 1&2 and 3&4 in **Figure 2c** are plotted in **Figure 2d** and show that neighboring electrodes (1 vs. 2; 3 vs. 4) can have different response properties, with 3 vs. 4 being a particularly stark contrast. This suggests a high degree of spatial localization of the recorded signals. These results demonstrate the ability to resolve the tonotopic organization of multiple auditory cortical fields with very high-resolution using high-frequency signals of CSEPs.

**Figure 2a-b** suggest that auditory responses of CSEP components from the same electrode are similar. The plots in **Figure 2e-f** display average normalized FRAs across all channels with tuned responses in the Hγ and tMuA components, which were indeed similar. To quantify this, we first determined a boundary that separated the responsive portions of the FRA from the unresponsive portions (e.g., white lines in **Fig. 2e-f**, see Methods). The average FRA-boundaries for different frequency components across all tuned channels (β = 260, γ = 292, Hγ = 333, uHγ = 302, tMuA = 113) are displayed in **Figure 2g** (mean ± s.e.), and were highly overlapping. We found that median best (auditory) frequencies (BF, extracted from FRA) were ∼8.5kHz (**Fig. 2h**), and there was a mild effect of CSEP component on BF (Kruskal-Wallis, df = 4, χ^2^ = 28, P < 0.001). The range of values in our data set (interquartile ranges, **Fig. 2h**) makes the functional relevance of the marginal statistical significance for best frequency questionable. Indeed, the best frequencies extracted from different components at an electrode were highly correlated (R, median = 0.89; range = [0.8 0.93], P < 10^−5^ for all). This indicates that different high-frequency CSEP components are generated by neurons with very similar tuning properties, perhaps from the same cortical column. Additionally, the median (auditory) bandwidths (BW, extracted from FRA as full-width at half-max) were ∼1.5 octaves (**Fig. 2i**) and there was a robust effect of CSEP component on BW (Kruskal-Wallis, df = 4, χ^2^ = 116, P < 10^−22^). Thus, there was a systematic decrease in bandwidth with increasing CSEP component in the electrical potential (**Fig. 2i**), perhaps reflecting greater spatial spread of the lower frequency signals.

### CSEPs are anisotropically localized to a cortical column

The spatial spread of electrical signals in the brain is of great interest, both for its importance in interpreting recorded electrical potentials and for its practical implications for sensor design. In principle, two primary factors that contribute to the spatial spread of correlations in electrophysiology recordings are diffusion due to the electrical properties of the tissue, and the spatial organization of neuronal function (e.g., tonotopy). If diffusion is the main determinant, then the spatial spread is expected to be isotropic. However, given the tonotopic organization of rat auditory cortex (**Fig. 2c**), we hypothesized stronger correlations in the isotonic dimension (DV) versus the heterotonic dimension (AP). Furthermore, because tonotopy arises from the organization of cortical columns which contain neurons with similar tuning properties^21,23^, we hypothesized that the CSEP signals would be tightly localized the approximate diameter of a column (200-500 μm)^21–23^.

To assess spatial spread, for each CSEP frequency component, we fit a general linear model to single-trial tone responses for each target electrode as a function of the responses at all other electrodes. Our analysis method, based on sparse general linear models, largely mitigates potential confounds due to pairwise chaining of local correlations by factoring out the covariance matrix of the regressors. We fit the linear model using the UoI_Lasso_ algorithm^33^ (see Methods) which provides accurate estimates of parameters values (R^2^>0.9 for all fits). For each target electrode, the parameter values from the fit model (weightings on responses at other electrodes) were normalized to their maximum value across frequency to ease comparisons across frequencies and electrodes. **Figure 3a-b** display the spatial distributions of median fit parameters as a function of location relative to the target electrode (demarcated as ‘X’) for all frequency tuned electrodes in the Hγ and tMuA components. For both of these signals, we found that model parameters were extremely localized in both AP and DV directions (values quickly go to zero), and exhibited a marked anisotropy, with larger values DV than AP. Also, parameters were larger for Hγ than for tMuA (grey scale), indicating that less tMuA variance could be explained by surrounding electrodes, and thus suggesting a more localized signal.

**Figure 3.**
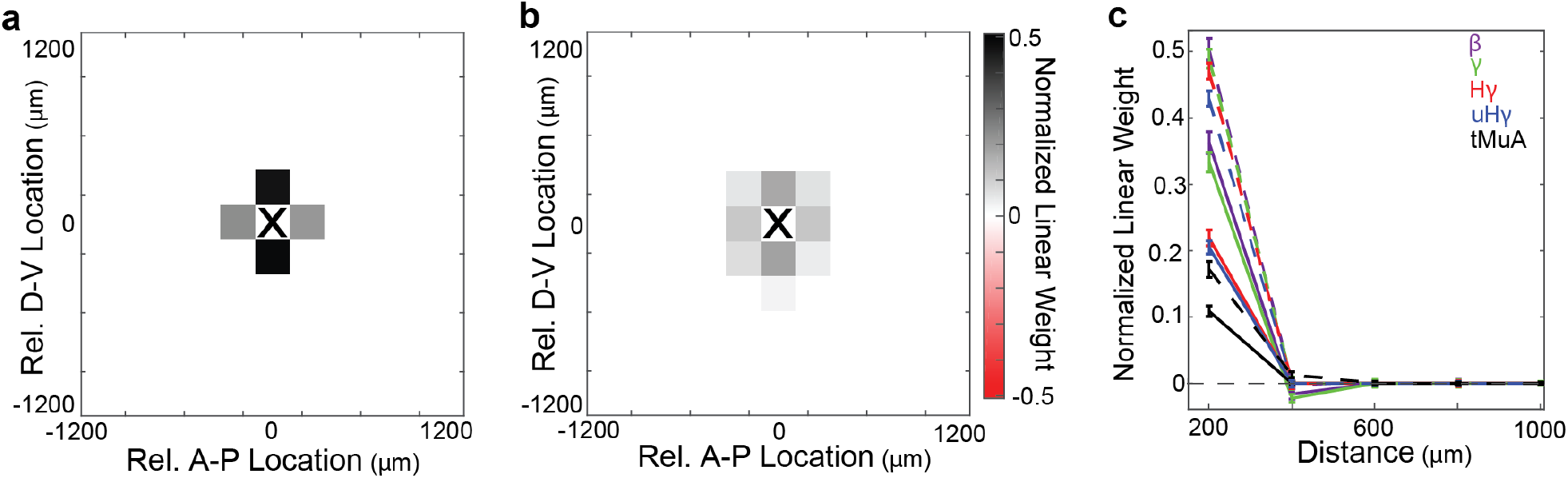
CSEPs are anisotropically localized to a cortical column. **a**. Spatial distribution of weights from a regularized linear model of Hγ responses during the tone stimuli as a function of the other electrodes on the grid. Locations are all relative to the electrode used as the dependent variable in linear regression. Values are median across all N = 333 tuned (in Hγ) electrodes. **b**. Spatial distribution of normalized weights for tMuA. Values are median across all N = 113 tuned (in tMuA) electrodes. **c**. Median ± s.d. of normalized linear weights across all electrodes as a function of distance in the AP (solid lines) and DV (dashed lines) dimensions along the grid. Different frequency bands are demarcated with colors.

We summarized the results of this analysis for the β through tMuA components (**Fig. 3c**, colors, N’s: β = 260, γ = 292, Hγ= 333, uHγ = 302, tMuA = 113) by plotting the median model parameters as function of distance in the AP (solid lines) and DV (dashed lines). Across frequency components, we found that ∼70% of the parameter magnitudes were concentrated at ±200 μm (4/143 grid locations with non-zero values, ∼3%), indicating that the vast majority of explanatory variation was localized immediately surrounding the electrode. Furthermore, the parameter values were significantly greater in the dorsal-ventral than the anterior-posterior direction for all CSEP components (Wilcoxon Sign Rank Test, P < 10^−4^ - 10^−28^). Within both the AP and DV directions, a significant effect of CSEP frequency component on parameter magnitude at 200 μm was observed, with lower frequencies having larger values (Kruskal-Wallis; df = 4; DV: χ^2^ = 35, P < 10^−6^; AP: χ^2^ = 51, P < 10^−12^). This indicates that lower frequencies have a greater spatial spread and is in line with lower frequencies having broader tuning (**Fig. 2**). A cortical column is ∼350 μm in diameter^22^. Thus, these results indicate that cortical surface electrical potentials are localized to single cortical columns, and that the degree of localization increases with increasing CSEP frequency. Furthermore, the localization is anisotropically distributed and aligned with tonotopic organization, indicating that differentiation of function across the cortical surface is the primary determinant of spatial correlations of evoked μECoG signals.

### Biophysical *in silico* cortical column reproduces *in vivo* observed μECoG response

The experimental results presented above demonstrate that evoked CSEPs were tightly localized along the cortical surface, approximately to a single cortical column in the rat. This suggests that the sources of the μECoG signal are mostly located within the column directly underneath the electrode. To investigate the laminar and cellular origin of sources within a column that generate CSEPs, we simulated a full-scale, biophysically detailed model of a cortical column where each neuron’s full morphology is represented by 100’s to 1000’s of connected cylindrical neuronal segments^28^. The column model and μECoG electrode is depicted in **Figure 4a**, where circles indicate the locations of (a subset of) neuronal somas (black: excitatory neurons; red: inhibitory neurons). Stimulus-evoked input to the column is provided by activating (with Poisson spike trains) thalamocortical synapses located throughout the column according to the distribution shown in **Figure 4b**, which also displays the cortical layers.

**Figure 4.**
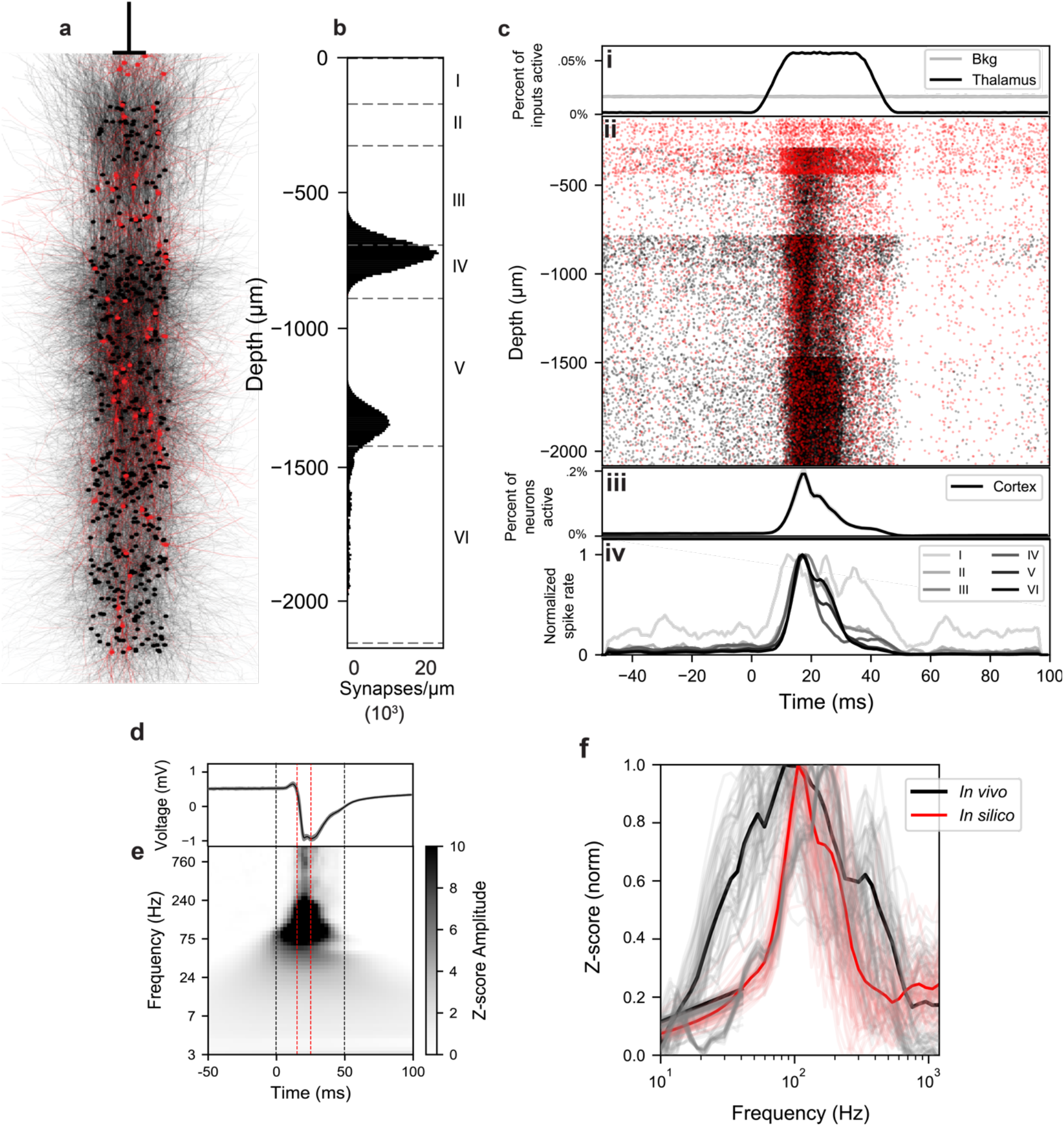
Biophysical *in silico* cortical column reproduces *in vivo* observed μECoG response. **a**. Rendering of a random sub-selection of 626 neurons in the simulated column (about 2% of the total). Black: excitatory neurons; red: inhibitory. Circles represent somas, lines represent dendritic structures. The position of the simulated μECoG electrode relative to the column is shown above. **b**. Distribution of synapses from the thalamus along the depth axis of the simulated cortical column. **c**. Data from one simulated stimulation and pre/post-stimulus silence **i**. Population spike rate of thalamic and background cortical spike trains activating synapses in the column. **ii**. Spike raster of all neurons in the column vs. soma depth (y-axis). Note that differences in raster density in part reflect differences in neuron density across cortical layers. **iii**. Population spiking (fraction of neurons spiking in 1 ms) of biophysically detailed cortical neurons. **iv**. Cell-averaged spike rate of biophysically detailed neurons in each layer. Darker shades indicate deeper layers. **d**. CSEP computed by the Line Source Approximation from all neurons in the column during a 150 ms window centered around the 50 ms “tone pip” stimulation. **e**. Spectrogram of the CSEP in panel **d**, z-scored to baseline **f**. Frequency content of CSEP during 10 ms centered at the response peak (indicated with dotted red lines in panels **d** and **e**), z-scored to baseline. Individual electrode averages from experimental results are in grey, black is grand average. Individual stimulus presentations from simulations are in pink, red is grand average. All traces are normalized to their respective maxima.

An example of the column’s activity is displayed in **Figure 4c**. The biophysical neurons in the column received thalamic input in the form of Poisson spike trains that were modulated in time to emulate our tone stimulus (**Fig.4c.i**, black) and background Poisson spike trains (**Fig.4c.i**, grey) that were not modulated by the stimulus. In **Figure 4c.ii** we show the spike times (black: excitatory neurons; red: inhibitory neurons) in response to one presentation of the input stimulus (**Fig.4c.i**). Neurons are arranged by depth below the surface, which allows us to visualize the laminar boundaries as sharp changes in the density of firing reflecting different cell densities across layers. The fraction of neurons in the column firing action potentials is displayed in **Figure 4c.iii** as a function of time. The time-to-peak of about 15-20 ms in most layers (**Fig. 4c.iv**), as well as the following period of slightly elevated activity until stimulus offset, are both consistent with the *in vivo* recordings^25,34^.

The biophysical model produces CSEPs (**Fig. 4d-f**) consistent with the high-frequency transient onset response observed *in vivo*. We computed the electrical potential at the cortical surface of the simulated column using the line source approximation^35^, and processed the simulated data identically to the experimental data. Average (mean ± s.e., N = 60 stimulus presentations) raw evoked cortical surface electrical potential from the model is plotted in **Figure 4d**, while the spectral content of the simulated CSEP is shown in **Figure 4e** (dashed black lines: stimulus; dashed red line: 10ms window around the peak high-gamma response). The extracted z-scored (max normalized) response as a function of frequency for the simulated CSEP is shown **Figure 4f**, as well as the experimental data (black). There is striking agreement in the high-frequency content of CSEPs collected experimentally and CSEPs generated by the simulations. Both experimental and simulated CSEPs exhibit a peak frequency of ∼100 Hz, and the spread of the signal around the peaks are overlapping. Thus, biophysical simulations accurately recreate key aspects of experimentally acquired data, indicating that they are a good forward model of CSEP generation.

### *In silico* cortical column predicts experimentally observed relationship between response magnitude and frequency

The model makes testable predictions regarding the relationship between the magnitude and frequency content of CSEP responses. In the simulations, we varied the magnitude of the excitatory input to the network by increasing the mean firing rate of the thalamic spike trains during the stimulus. Analogously, in the experiments, we monitored the evoked CSEP in response to varying sound amplitudes at each electrode’s best frequency.

For the simulations, **Figure 5a** displays the average normalized evoked response to stimulation of different amplitudes as functions of frequency (input magnitude given by color saturation, indicated by inset color bar). We observed that the magnitude of CSEP response depended monotonically on input magnitude. More interestingly, we found that as the magnitude of the response increased, so did the frequency content of that response. This can be seen as a sweep towards the upper right of the individual traces as input magnitude increases. We quantified the relationship between response magnitude (z-scored response between 10 and 200 Hz) and the frequency content (frequency at peak response between 10 and 200 Hz). The pink-to-red squares in **Figure 5c** display the normalized maximum response magnitude vs. normalized frequency at maximum response for varying input amplitudes (color saturation demarcates magnitude of input, circled square demarcates input used in Figures 4, 6, 7). Intuitively, these effects were mediated by an increase in the population mean firing rate and spike synchrony resulting from increased input spike rate.

**Figure 5.**
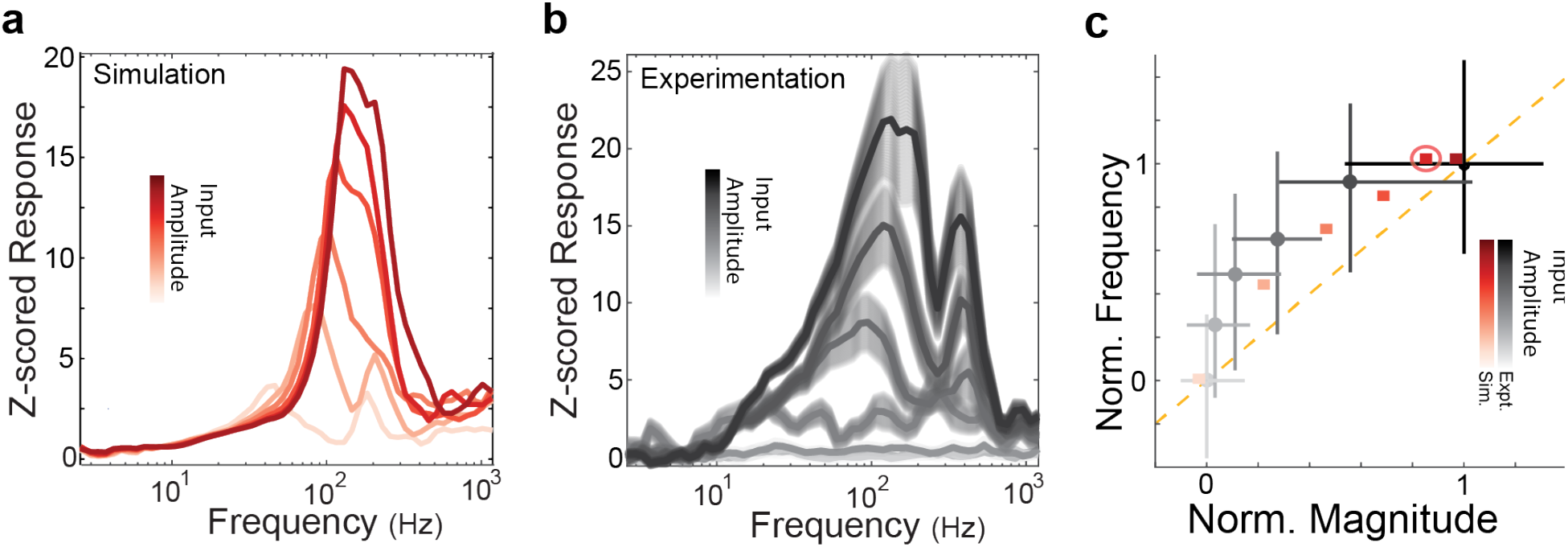
*In silico* cortical column predicts experimentally observed relationship between response magnitude and frequency. **a**. Average z-score as a function of frequency in eight simulations with variable input amplitude. **b**. Average z-score as a function of frequency in the experimental data for six different stimulus amplitudes. **c**. Normalized response magnitude vs. normalized response frequency for experimental data (black, mean ± s.d.) and for simulations (red). Each data point corresponds to the response frequency and magnitude associated with a distinct input magnitude (response magnitude increases monotonically with input magnitude). Circled point indicates the input magnitude used in Figures 4,6,7. Orange dashed line is unity.

**Figure 6.**
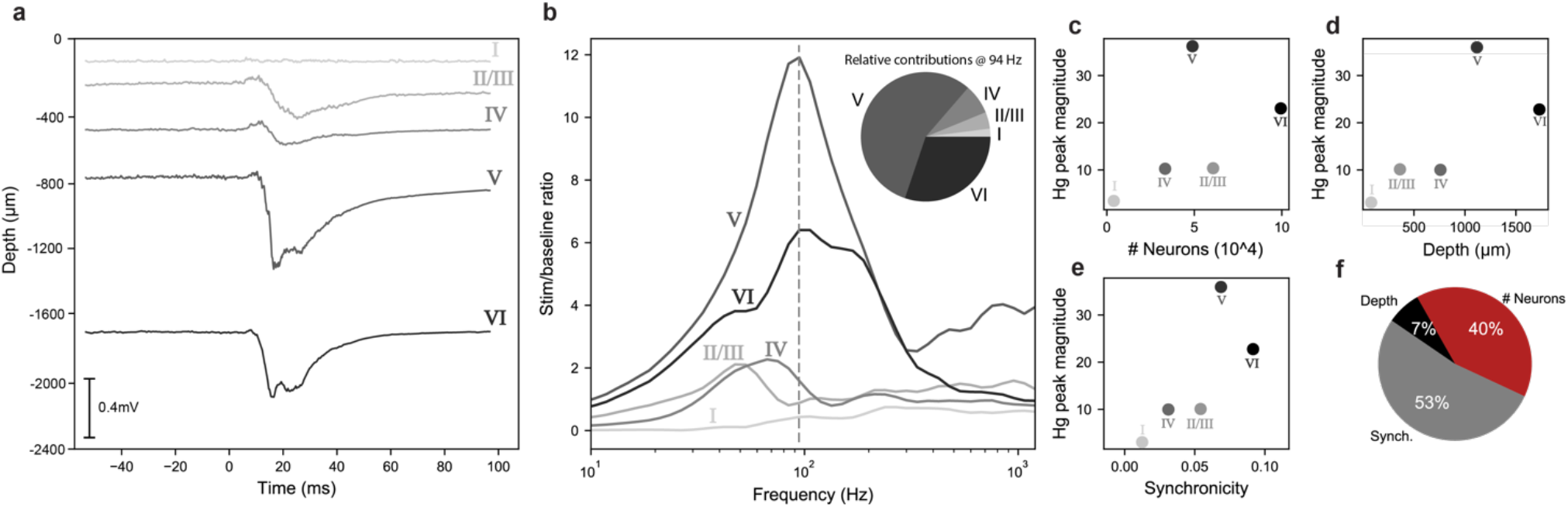
Evoked μECoG responses originate in infragranular layers. **a**. Contributions to the simulated CSEP from anatomical layers. Top-to-bottom: cortical layers I through VI. The sum of these contributions is the total CSEP. **b**. Frequency content of the laminar contributions during stimulus peak. Layer V and VI contributions dominate the high-gamma peak. **c**. Magnitude at peak frequency of each cortical layer’s CSEP contribution vs. number of neurons in the layer. **d**. Magnitude at peak frequency of each cortical layer’s CSEP contribution vs. average distance of cell bodies in the layer from the recording electrode. **e**. Magnitude at peak frequency of each cortical layer’s CSEP contribution vs. synchronicity of somatic membrane potentials averaged over all pairs of neurons in the layer. **f**. Pie chart showing the relative importance of these three factors in a linear model of the high-gamma peak contribution magnitudes of anatomical layers.

**Figure 7.**
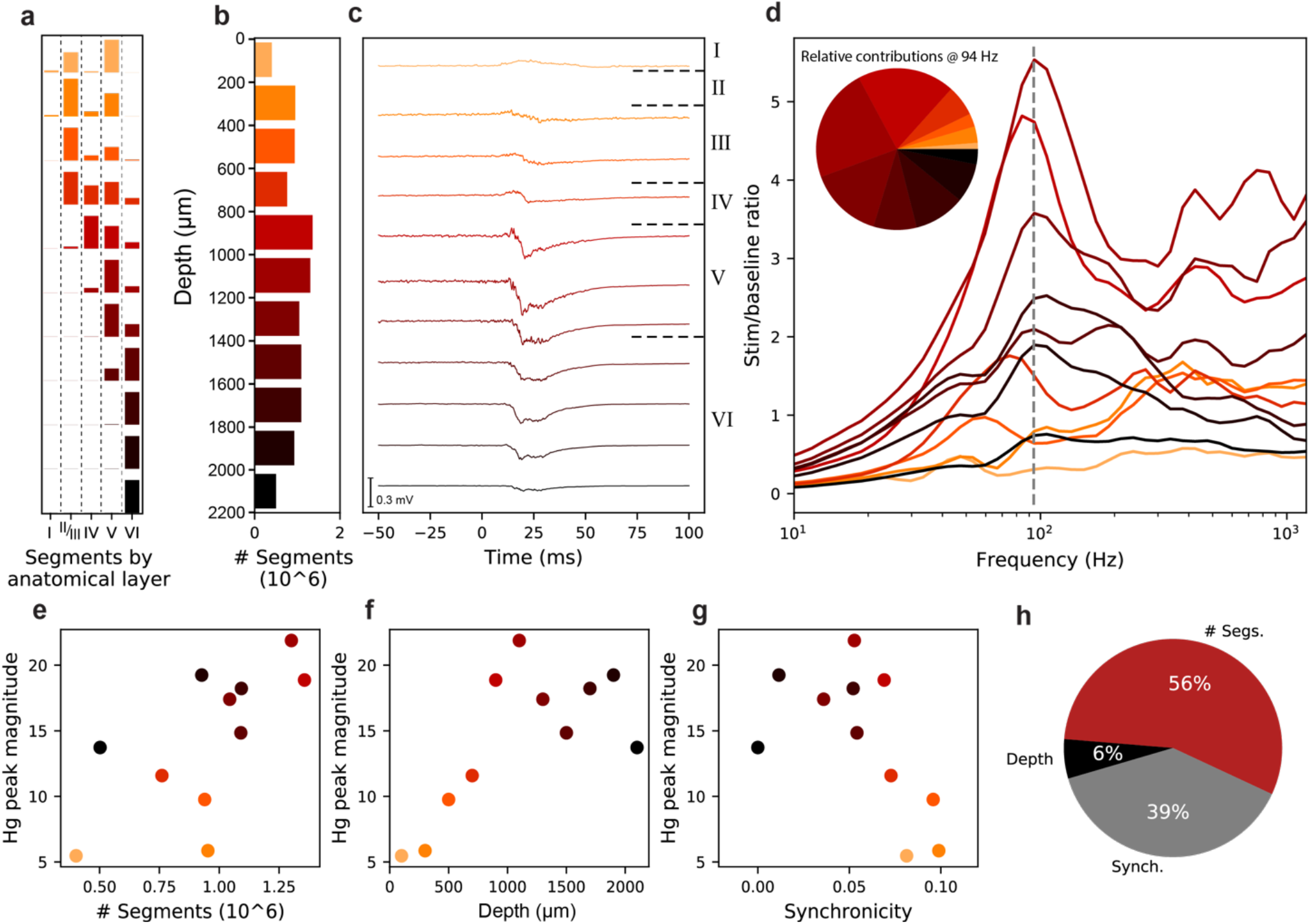
Evoked μECoG responses originate in sources 800-1400 μm below the surface. **a**. Proportional breakdown of segments by anatomical layer. Most slices contain segments from neurons in multiple cortical layers. Bars represent proportion of total segments in the slice, different slices not to scale. **b**. Total number of simulated neuronal segments in each 200 μm axial slice of the column. **c**. Contributions to the CSEP from 200 μm slices, organized by depth (top: cortical surface). The sum of these contributions is the total CSEP shown in Figure 4b. **d**. Frequency content of the slice contributions during stimulus peak, colored by slice depth. Slices containing somas of layer V neurons dominate the high-gamma peak. **e**. Magnitude at peak frequency of each slice’s CSEP contribution vs. number of neuronal segments in the slice. **f**. Magnitude at peak frequency of each slice’s CSEP contribution vs. average distance of segments in the slice from the recording electrode. **g**. Magnitude at peak frequency of each slice’s CSEP contribution vs. average synchronicity in the slice. **h**. Pie chart showing the relative importance of the three factors in our linear model of the slices’ high-gamma peak contribution magnitudes.

Next, we sought to determine if this relationship between magnitude and frequency existed in the experimental data. **Figure 5b** displays the z-scored CSEP at an example electrode as a function of frequency in response to the BF stimulus presented at different amplitudes (see inset color bar). Similar to **Figure 5a**, we observed a sweep towards the upper right of the individual traces with increasing input amplitude (**Fig. 5b**). For frequency tuned electrodes, we calculated the same quantities (maximum response magnitude and frequency at maximum response) as a function of the amplitude of auditory input at the electrode’s best frequency in the tone stimuli (**Fig. 5c**; grey-to-black circles, mean ± s.d., N ∈ [206 299]). As in the simulations, we observed that increasing the input magnitude resulted in an increase in both the magnitude of the peak response and frequency at the peak response. Further, there is a striking correspondence in the curvature of response frequency vs. response magnitude plots derived from experimental and simulation data (**Fig. 5c**). These results demonstrate a prediction made by the model that was confirmed by a novel experimental finding.

### Evoked μECoG responses originate in infragranular layers

We next utilized the model to understand the spatial distribution of the generating sources of the CSEP. A key feature of the biophysical model is that CSEP calculation is separate from the numerical simulation of the neurons in the column, enabling us to calculate CSEPs from arbitrary samples of neuronal segments in the column without perturbing the activity at all. We first examined the contributions to the CSEP from cortical layers by computing each layer’s contribution to the CSEP individually (see Methods).

**Figure 6a** plots the raw evoked CSEP (scale bar in inset) as a function of time for each layer, and indicates the average depth of neuronal somas for the layers. Surprisingly, we found that layers V and VI produce the largest evoked potentials, despite being the furthest away, while neurons in superficial layers contribute very little of the total CSEP. **Figure 6b** shows the frequency content of each contribution during a 10ms window surrounding the response peak. The inset shows the relative magnitudes of the layers contributions in the band centered at 94 Hz, the apex of the high-gamma peak, shown as a dotted vertical line in the main panel. As with the raw evoked potential, we found that infragranular layers also contribute most to the high-gamma component of ECoG responses: 51% from layer V, 35% from layer VI, and the remaining 14% coming from layers I-IV. To further probe the dependence of the high-gamma component on individual layers, we also performed lesion studies examining the CSEP produced when activity in one cortical layer is excluded, which showed that only layer V lesions were capable of flattening the high-gamma peak (**SFig.1**).

The results above appear counter-intuitive when the contribution of sources is viewed only as a function of distance. However, in addition to distance, the number of sources and their correlations are additional biophysical factors that dictate the contribution of neuronal populations to a distally recorded signal^20^. *A priori*, the relative importance of these factors to determining laminar contributions to evoked CSEPs in a full-scale cortical column model is not clear. Thus, we plotted each layers’ peak high-gamma responses as a function of the number of simulated neurons in **(Fig. 6c)**, the average distance of somas in each layer from the recording electrode **(Fig. 6d)**, and the synchronicity between somatic membrane potentials in each layer during the stimulus **(Fig. 6e)**. We note that in **Figure 6d** the peak high-gamma response vs. depth shows a positive slope, contrary to the physical principle that individual neurons further from the electrode will contribute less to the signal. However, as is evident from these plots, there are correlations between depth and the other variables. For example, deeper layers tend to contain more neurons. Thus, we fit a regularized linear model to predict peak high-gamma magnitude across layers as a function of depth, number of segments, and synchronicity of neuronal somas simultaneously, which fit the data well (R^2^ = 0.98). The relative magnitudes of the fit coefficients are plotted in **Figure 6f**, which shows that the number of segments and between cell synchronicity are the dominant factor that determine source contributions to CSEPs, while depth was a minor factor. To determine how robust these results were to baseline normalization, we performed the same analysis with a different normalization procedure and found very similar results (**SFig. 2**). Thus, infragranular layers contribute ∼86% of evoked CSEP responses because of their increased number of neurons and increased synchronicity.

### Evoked μECoG responses originate in sources 800-1400 μm below the surface

The previous results indicate that layers V and VI are the dominant sources to evoked CSEPs. However, due to the large, extended morphology of some neurons relative to the column depth, knowledge of the largest contributing anatomical layers does not necessarily imply precise knowledge of the spatial distribution of segments generating CSEPs. For example, the apical tufts of many layer V pyramidal neurons reach into layer I. Thus, we next isolate contributions to the CSEP from 200 μm axial slices of the column.

Most slices contain segments from neurons in more than one layer, and a given neuron can contribute to more than one slice. The breakdown of segments in each slice by anatomical layer is shown in **Figure 7a**, where each color represents one slice. For each slice, five bars are shown displaying the number of segments in that slice belonging to neurons in the five cortical layers. For example, the top slice is dominated by segments from layer V neurons (**Fig. 7a**, 4^th^ column). The total number of neuronal segments in each slice is shown in **Figure 7b**, which makes clear that the slices between 800-1200 μm have the most segments. **Figure 7c** shows CSEPs calculated only from segments in the slices as a function of depth (CSEP scale bar is inset). The largest contributors to the evoked responses are the slices located from 800-1400 μm below the surface, i.e., in layer V. We extracted a 10 ms window around the peak of the CSEP response and analyzed the frequency content of each slice’s contribution within that window. The results are shown in **Figure 7d**. The inset shows the relative magnitudes of the slices’ contributions at 94 Hz, the apex of the high-gamma peak, shown as a dotted vertical line. Here we see that the slices spanning 800-1400 μm are also the ones contributing most to the high-gamma peak (56% total), which is where layer V somas are located. Thus, this analysis demonstrates that layer V somas are the major generating source of evoked ECoG signals.

As with the layer contributions, we sought to ascertain the relative importance of the number of segments in the slice (**Fig. 7e**), the depth of the slice below the surface (**Fig. 7f**), and the synchronicity of membrane potentials of segments within the slice (**Fig. 7g**) in determining the high-gamma peak contribution magnitude. The results of a regularized linear regression predicting high-gamma peak from those parameters (R^2^ = 0.91) are shown in **Figure 7h**. As with the layer contributions, very similar results were observed using a different normalization procedure (**SFig. 3**). Similar to the layer contributions, we find that the number of segments and the synchronicity are the most important factors determining the magnitude of a slice’s contribution to the CSEP.

## Discussion

We found that evoked cortical surface electrical potentials had a strongly non-monotonic frequency structure, with a large peak in the 65-170 Hz range (Hγ). In the brain, the most prominent biophysical sources with energy in that frequency range are action potentials of pyramidal neurons^3^. In experimental data, we quantitively demonstrated that evoked CSEPs were localized to a cortical column. Full-scale biophysical simulations of a cortical column reproduce the experimentally observed evoked spectrum and indicate that evoked μECoG high-gamma is generated by infragranular neurons.

The spatial spread of electrical potentials (i.e., LFPs) is of long-standing debate and great interest both for basic neuroscience as well as practical applications to sensor design and brain-machine interfaces. We demonstrate a tight, anisotropic localization of evoked CSEPs to ≤±200 μm on the surface, with higher frequency components exhibiting greater spatial localization. Indeed, we observed examples of neighboring electrodes (in different auditory areas) with very different tuning properties. A column is ∼300 μm in diameter (range: 200-500 μm) ^22^. Thus, our results quantitatively demonstrate that CSEPs are localizable to individual cortical columns.

Generally speaking, our results indicate a much more localized signal than has been directly quantified in previous (μ)ECoG studies^16,18,36,37^, though there have been qualitative descriptions of tight localization^38^. Most studies of ECoG that directly quantify evoked signal spread do so by calculating the pairwise correlation between electrodes, and examine the decay of correlation with distance^16,18^. However, such pairwise analyses potentially confound long-range correlations due to direct interactions with correlations due to chaining of local (pairwise) interactions. The analysis methods used here, based on sparse general linear models, largely mitigates such confounds by factoring out the covariance matrix of the regressors. Our results corroborate and extend a study of the spatial spread of low-frequency electrical potentials inside the cortex^17^. However, in contrast to that report, we demonstrate that higher frequencies are more spatially localized and have narrower stimulus tuning. We note that our estimate of spatial spread should be considered an upper-bound, as it is at the limit of the inter-electrode spacing (which is still half that of Utah arrays).

We observed a large anisotropy in the spatial spread of correlations in evoked CSEPs. The anisotropic localization had greater values in the dorsal-ventral direction than the anterior-posterior direction. The anisotropic spread thus reflects the functional organization (tonotopy) of the underlying cortical tissue^21,31,32^. This indicates that the major determinant of the spatial scale of correlations in stimulus evoked high-frequency electrical potentials across the surface is not the passive propagation of signals through the cortical tissue (which is expected to be isotropic^26^), but instead is correlations in function. As cortical columns are the basis of the functional organization of cortex^22^, the anisotropy corroborates localization to a cortical column. Together, our results imply that maximization of ECoG for basic neuroscience and brain-machine interfaces requires small electrodes with very tight electrode spacing and electronics with good signal-to-noise at high frequencies. In particular, ECoG grids for human BMI should be matched to the spatial resolution of functional differentiation of the underlying cortex, and subsampled relative to this to ensure robustness to electrode loss over time.

Like all electrophysiological methods, ECoG records compound electrical potentials (i.e., LFPs) that reflect a weighted linear superposition of all electrical sources in the brain^3,20,26^. Several studies have examined the relationship between intracortical spiking activity (both single-unit and multi-unit activity, MuA) and the frequency content of simultaneously recorded intracortical LFPs, but report divergent results. For example, Steinschneider et al.^39^, characterized the spectral content of A1 evoked LFPs and the relationship to MuA in non-human primates. Several studies have suggested that high-frequency content of intracortical LFPs (‘broad-band activity’) is directly modulated by neuronal spiking activity^40,41^. Ray & Maunsell^42^ showed that evoked intracortical gamma band activity can be dissociated from high-gamma band activity, while a report from Leszczynski^43^ suggests that broadband activity can be dissociated from MuA in different layers. However, these studies have primarily focused on intracortical recordings. Thus, little is known about the relative contributions of the superposed neuronal sources at the cortical surface.

Our study addresses this gap with a full-scale, biophysically detailed model^28^ of cortical sources and forward model of their superposition at the surface. This model allowed us to interrogate the laminar and cellular contributions to CSEPs using manipulations that would not be possible experimentally, such as isolating a single anatomical layer or slice of cortex without disturbing the activity of the column. The model produces stimulus-evoked responses with high-frequency content strikingly similar to that observed experimentally. In both simulations and experiments, the largest magnitude responses were observed in the high-gamma range, peaking at ∼100 Hz in both cases. The simulation predicted that as the magnitude of the input increases, there should be concomitant increases in both the frequency and amplitude of the evoked response with a concave shape. This prediction was validated with a novel observation in the experimental data. Previous efforts to understand CSEPs and intracortical LFPs produced by large populations of neurons use simplified neuronal models^14,44^, or simplified network models that omit several cell types or even entire cortical layers^45^. The Blue Brain Project model^28^, used here, represents the state of the art in biophysically detailed simulation of neurons and cortical columns. Together, our results indicate that, for the first time, a computational model accurately captures the biophysical processes giving rise to the evoked ECoG response observed in the data.

The biophysical model demonstrates that the intuition that CSEP signals must be generated primarily in superficial layers is incorrect. Instead, our analysis implicates neurons in cortical layer V as the primary source of the signal recorded at the surface, with layer VI also contributing substantially. Similarly, analysis of contributions to the CSEP by depth showed that slices of the column containing layer V somas produce most of the signal observable at the surface. Subsequent analysis found that the density (number) and synchrony of neurons to be more important than depth in determining a population’s contribution to the surface signal. While layers II/III/IV are closer to the recording electrode than layers V/VI, they have fewer neurons^2,28^ and reduced synchrony^46,47^. Layers V/VI are composed of predominately excitatory pyramidal cells^28,47^, and pyramidal neuron action potentials contribute most to the high-frequencies examined here^3^. Thus, we conclude that evoked high-gamma at the surface is a biomarker of layer V/VI pyramidal neuron firing rates.

The simulation results suggest a word of caution for the interpretation of multi-unit activity (and LFPs more broadly) recorded both at the surface and intracortically^9,48,49^. In particular, we found that activity in the multi-unit activity range (500-1000Hz) at the surface was predominantly generated by neurons in layer V, sources which are very distant (1-1.5mm) from the recording electrode. Hence, previous reports of ‘single-unit’ recording from the cortical surface, as well as intracortically recorded multi-unit activity (e.g., laminar polytrodes, Utah arrays, etc.) may contain contributions from distal, but numerous and synchronous, neurons. The finding that evoked ECoG high-gamma is primarily generated by neurons in layer V provides a potential explanation of the robust tuning to exogenous variables found here and elsewhere (e.g., auditory stimuli, vocal tract articulators^6^, etc.,). In particular, neurons in layer V have previously been found to have sharper tuning curves than neurons in layer II/III^25,34,50^. Thus, while cortical surface electrical stimulation may activate broadly connected neurons in layer II/III^50,51^, recordings from the surface can reflect the finely tuned responses and precise projections of neurons in layer V^34,50^. The homogeneity of cortical columns within an area^22^ and the linear properties of the cortical tissue suggests that the results here derived from μECoG will extrapolate to the larger electrodes used in the clinic.

In summary, our results indicate that evoked ECoG high-gamma responses are primarily generated by the population spike rate of pyramidal neurons in layer V/VI of single cortical columns. Together, these results highlight the possibility of understanding how microscopic sources (specific neuronal populations) produce mesoscale signals (i.e., ECoG). For example, in some cases we observed a pronounced secondary peak at ∼375Hz in the experimental data (**Fig. 5b**); this novel spectral component has not been previously characterized, and was not reproduced by the single column simulation. We conjecture that this novel spectral component may reflect the activity of neurons in layer II/III in response to input from adjacent cortical columns (which were not explicitly modeled here). More broadly, we propose that different high-frequency components of ECoG signals (e.g., high-gamma, ultra-high-gamma, multi-unit activity, etc.,) reflect spiking activity of neurons in different cortical layers. As neurons in different layers are thought to perform distinct computations^47^, this proposition implies that different components of ECoG signals could be biomarkers for these computations. Experimentally, this could be addressed by recording ECoG with laminar polytrodes combined with layer/cell-type specific optogenetic perturbations^52,53,54,55^ analyzed with current source density estimation^56^. However, the highly interconnected nature of neurons in a cortical column makes independent and isolated control of only one neuronal population challenging at best. Further, our results indicate that considering properties of the population (e.g., synchrony and number of sources) are essential. Thus, biophysically detailed simulations of evoked CSEPs combined with experiments will remain a critical tool for understanding the origins of experimentally observed ECoG signals^20,26–28^.

## Author Contributions

K.E.B. designed the μECoG grid with input from P.L.; M.M.M and P.L. developed the μECoG grid. K.E.B. and M.E.D. conducted the rat experiments. K.E.B. and M.E.D. analyzed the data. V.L.B. implemented the simulations and analyzed the data with input from K.E.B.; K.E.B., M.E.D., and V.L.B. wrote the paper with feedback from P.L.; K.E.B. conceived of and supervised the project.

## Acknowledgements

LBNL-internal LDRD “Neural Systems and Data Science Lab” and NINDS R01(RNS118648A) (KEB); Marco Microelectronics Advanced Research Corporation 2009-BT-2052 and NSF grants EFRI 1240380 and DBI-1152658 (MMM).

## Methods

### Key Resource Table

(to be included upon acceptance)

### Contact for Reagent and Resource Sharing

Further information and requests for resources should be directed to and will be fulfilled by the Lead Contact, Kristofer E. Bouchard.

### Experimental Model and Subject Details

Data from four rats (female Sprague Dawley) were used in this study. All animal procedures were performed in accordance with established animal care protocols approved by the Lawrence Berkeley National Laboratory, Institutional Animal Care and Use Committees.

### Method Details

#### Electrophysiological recordings

All neural data were recorded with a multi-channel amplifier optically connected to a digital signal processor (Tucker-Davis Technologies [TDT], Alachua, FL). Signals were acquired at 12 kHz and lowpass filtered to the Nyquist frequency (6 kHz).

#### Rodent Preparations

We performed experiments in four anesthetized female Sprague Dawley rats. Animals were given a 1 mg/kg subcutaneous (s.q.) injection of Dexamethasone the night before a procedure to reduce cerebral edema. An anesthetic state was induced with an inductive dose of ketamine (95 mg/kg i.p.) and xylazine (10 mg/kg i.p.). Anesthetic state was assessed using toe pinch reflex and monitoring respiration rate. Additional doses of ketamine (55 mg/kg i.p) and xylazine (5 mg/kg i.p.) were administered as needed to maintain a negative reflex and a regular reduced respiration rate. Respiration was supported with a preoperative subcutaneous injection of atropine (0.2 mg/kg), and a perioperative nose cone supplying .8L/min of O_2_. A water heating bed provided thermostatic regulation. To prevent dehydration over the 10-hour surgery and recording session, subcutaneous saline injections (1 ml/kg) were provided every 3 hours. Once anesthetized the rodent was affixed to a snout stereotax without earbars.

After stable anesthetic state was achieved, an incision was made along the sagittal midline. All the soft tissue on top of the skull was removed to reveal the lambda and bregma fissures. Two 1mm burr holes were drilled over non-auditory cortical areas–one between the lambda and bregma on the left hemisphere, and another anterior to bregma on the right hemisphere. These serve to reduce intracranial pressure and provide a reference for electrophysiological recordings. The right masseter muscle was then transected to uncover the portion of cranium lying over the right auditory cortex. Using a 1mm diamond tapered round Stryker dental drill, a craniotomy was performed to expose the cortex.

### Recording Devices

Rodent electrophysiological recordings were made with custom designed 128-channel μECoG grids (initially fabricated in-house, purchased at the time through Cortera Neurotechnologies, Inc., Berkeley, CA). These arrays were placed over the primary auditory cortex as identified by anatomical and physiological properties. For the rat experiments, the dura was surgically removed and the μECoG grid was placed directly on the pial surface. The μECoG grids were placed over the primary auditory cortex and grounded via a silver wire inserted into a non-auditory cortical area in the contra-lateral hemisphere, which also served as the reference. Each contact on the grid had an impedance of 30 ± 10 kΩ after electroplating with platinum black (measured in 1x phosphate-buffered saline at 1kHz), and had an exposed diameter of 40 μm, with a 200 μm inter-electrode pitch.

### Auditory Stimuli

Pure tone pips (50ms in duration) were played through a DVD player (Sony), attached to a TDT amplifier and played through an electrostatic speaker located near the left ear of the animal (80dB SPL, A-scale). The frequency and amplitude of the tone pips were parametrically varied. The frequencies spanned 6 octaves (500 Hz to 32 kHz) in 30 increments and 8 attenuations from 0 to -70dB (0dB attenuation at 80 dB SPL). Each frequency-attenuation combination was presented 25 times in pseudorandom order with 250 ms of inter-stimulus silence.

### Data Analysis

All analysis was performed using code written in Matlab (The Mathworks) or Python. All software is available upon request.

### Spectral analysis of cortical surface electrical potentials (CSEPs)

We calculated the spectrogram of the entire recorded electrical potential time series for each electrode from 4-1200 Hz (54 bins) using a constant-Q wavelet transform^57^. Constant-Q refers to a time-frequency decomposition in which frequency bins are geometrically spaced and Q-factors (ratios of the center frequencies to band-widths) are equal for all bins. The non-causal component of displayed responses (e.g., **Figure 1d**) is due to the large bandwidth at lower frequencies of our constant-Q time-frequency transform.

The amplitude for each frequency bin in the neural signal was normalized relative to baseline statistics by z-scoring. The baseline statistics were computed from the periods of silence between each stimulus presentation (the 50 ms immediately following each stimulus presentation is excluded from the baseline period). Z-scoring largely removes the canonical ∼1/f^α^ falloff of power with frequency (characteristic of many natural signals), highlighting stimulus evoked changes. We note that this procedure is preferred to examining the residuals from a ‘fit and subtract’ methodology, given the potential issue of fitting power-laws especially at the extremes of the frequency range.

Responses to stimuli were taken as the average z-scored activity ±5ms around the peak response time after the onset of the auditory stimulus. Peak response time was calculated using the High Gamma (65-170 Hz) component of the signal, which was computed as the average of z-scores for frequency bands whose center frequency falls in that range. The shielding and grounding of our rodent experimental recording systems was sufficient to avoid any significant 60 Hz noise in the μECoG recordings. We determined z-scores for the canonical neural frequency bands by taking the mean z-scores across frequency bins in the corresponding frequency range; Beta (10-27 Hz), Gamma (30-57 Hz), High Gamma (65-170 Hz), Ultra-high Gamma (180-450 Hz), and the Multi-unit Activity Range (500-1100 Hz).

### Analysis of responses to pure tones

The frequency response area (FRA) depicts the modulation of a CSEP components response as a function of frequencies (*F*) and amplitudes (*A*). For each frequency-amplitude pair (*f,a*) in the stimulus set (see above, Auditory Stimuli), we identified the response peak, *y*_*f,a,i*_, as the maximum amplitude between 10 and 20ms after the onset of stimulus *i*. We took the mean z-scored response across *n* trials:

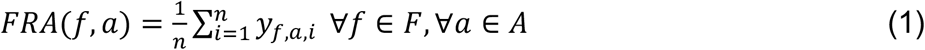

We estimated an FRA boundary to define a set of frequency-amplitude pairs that evoked a response. Intuitively, this is the stimulus-driven portion of the FRA, and corresponds with the part of the FRA that resides within the canonical ‘V’ shape. These boundaries were extracted using an approach identical to one used in a previous study characterizing rodent auditory responses^31^. A response-frequency function, *FRA*_*a*_(*f*), was computed by taking the mean of the FRA across all amplitudes, *n*_*a*_, and a response-amplitude function, *FRA*_*f*_(*a*), was computed by taking the mean of the FRA across frequencies, *n*_*f*_:

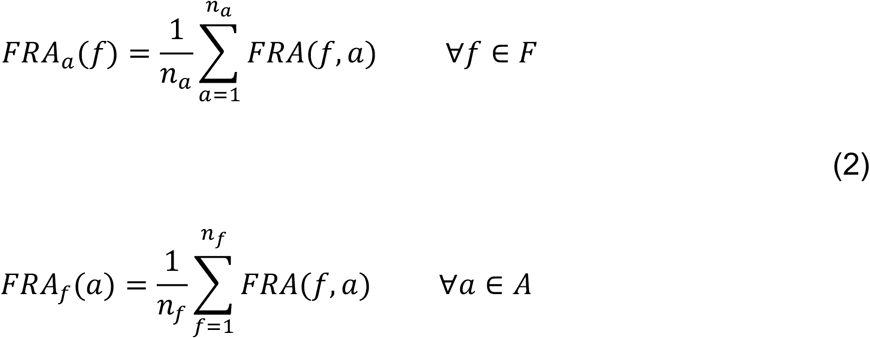

The *FRA*_*a*_(*f*) function defined the shape of the FRA boundary, and the inflection point of the *FRA*_*f*_(*a*) function, *a*_*b*_, was used to scale and shift that FRA boundary:

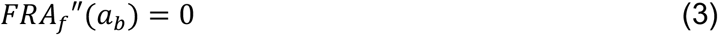

To fit the FRA boundary, *B*(*f*), we negated the response-frequency function, normalized its height, producing the normalized FRA, *normFRA*_*a*_(*f*), which was overlaid on the FRA. We constrain *normFRA*_*a*_(*f*^*^) to be 0, where f* is the value of f that minimizes *normFRA*_*a*_(*f*). The negated sound-frequency function was shifted via additive constant until its minimum coincided with, *a*_*b*_:

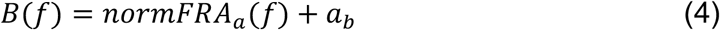

Portions of the FRA boundary that fell outside of the FRA were removed, resulting in the final FRA boundary. This method yielded FRA boundaries that closely resemble those curated manually.

We determined if a recording site was tuned using a permutation test. For a given FRA, we once again define *FRA*_*a*_(*F*) as mean of the FRA values across stimulus amplitudes. The standard deviation of the *FRA*_*a*_(*F*) function, evaluated on the data is a measure of a site’s tuning to a narrow band of frequencies. We compared the standard deviation (σ) of the original *FRA*_*a*_(*F*) with a null distribution *FRA*_*rand,a*_ consisting of 100 random shufflings of that FRA along both the frequency and amplitude axes. A channel is considered tuned if the standard deviation of its average stimulus frequency response, *FRA*_*a*_(*F*), exceeds the 95th percentile of a null distribution generated from 100 permutations of the FRA.

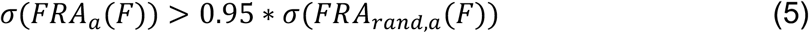

A tuned response exhibits a larger standard deviation across frequencies than an untuned (flat) *FRA*_*a*_(*F*). Randomly shuffling a tuned FRA removes the tuning structure and consequently reduces the resulting standard deviation. All subsequent tone analysis only included tuned sites.

We used the FRA boundary to further characterize the response properties of each recording site. We defined a best frequency (BF) as the weighted average of the FRA boundary, and calculated bandwidth (BW) as the full width of FRA boundary at half-maximum, *FWHM*:

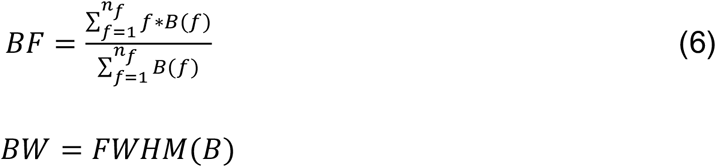

### Spatial Analysis of CSEPs

We assessed the spatial spread of CSEPs by fitting a general linear model. For a given CSEP component, the responses at an individual single electrode (*R*_*i*_) was modeled as a linear combination of responses at all other active electrodes (*R*_*j* ≠ *i*_), corrupted by independent and identically distributed (i.i.d.) Gaussian noise (*ε*):

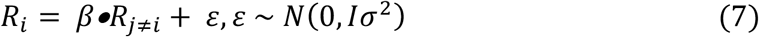

Where *I* is the identity matrix. For each frequency component separately, both the dependent (*R*_*i*_) and independent responses (*R*_*j* ≠ *i*_) are standardized to have a mean response of 0 and a variance of 1 (thus, there is no y-intercept term). This standardization enables comparison of weights across frequency component. After fitting, we arranged the estimated weights 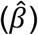 for each recording site (independent variables) relative to the position of the dependent variable (electrode i) on the μECoG grid, which allowed us to compare the relative spatial distribution of weights across all recording sites.

### The UoI_Lasso_ algorithm

As mentioned above, we used a novel statistical inference procedure (UoI_Lasso_^33^) to fit the general linear model in equation (7). While a detailed description of this algorithm is outside the scope of this manuscript, here we provide the motivation, outline the innovations, and summarize the main statistical result of these innovations. The interested reader is encouraged to see Bouchard, et al., 2017, for further details.

Generally speaking, in regression and classification, it is common to employ sparsity-inducing regularization to attempt to achieve simultaneously two related but quite different goals: to identify the features important for prediction (i.e., model selection) and to estimate the associated model parameters (i.e., model estimation). For example, the Lasso algorithm in linear regression uses L_1_-regularization to penalize the total magnitude of model parameters (‖*β*‖_1_), and this often results in feature compression by setting some parameters exactly to zero. Using the notation in equation (7) for a given ‘target’ electrode (*R*_*i*_), this corresponds to solving the constrained convex optimization problem:

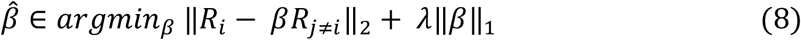

It is well known that this type of regularization implies a prior assumption about the distribution of the parameter (e.g., L_1_-regularization implicitly assumes a Laplacian prior distribution). However, strong sparsity-inducing regularization (i.e., large values of *λ*), which is common when there are many more potential features (p) than data samples (n) (i.e., the so-called small n/p regime) can severely hinder the *interpretation* of model parameters. For example, while sparsity may be achieved, incorrect features may be chosen and parameters estimates may be biased. In addition, it can impede model selection and estimation when the true model distribution deviates from the assumed distribution.

To overcome these and other issues, we have recently introduced a novel statistical-machine learning framework called Union of Intersections (UoI)^33^. Methods based on UoI perform model selection and model estimation through intersection and union operations, respectively, leading to enhanced model selection and estimation. Focusing on linear regression, the UoI_Lasso_ algorithm has three central innovations: (1) calculate model supports (S_l_) using an intersection operation over bootstrap resamples for a range of regularization parameters λ (increases in λ shrink all values of β towards 0), efficiently constructing a family of potential model supports {*S*: *S*_*l*_ ∈ *S*_*l*+*k*_ for k sufficiently large}; (2) use a novel form of *model averaging* in the union step to directly optimize prediction accuracy (this can be thought of as a hybrid of bagging and boosting); and (3) combine pure model selection using an intersection operation with model selection/estimation using a union operation in that order (which controls both false negatives and false positives in model selection). Together, these innovations lead to state-of-the-art selection (the selected parameters are a Union of Intersections, hence the name), estimation, and prediction accuracy. This is done without explicitly imposing a prior on the distribution of parameter values, and without formulating a non-convex optimization problem.

### Biophysically detailed simulation of cortical column

Using the NEURON simulation environment running on 64 nodes of Cori (Cray XC70) at NERSC, we implemented and successfully executed a biophysically detailed model of a cortical column and the CSEP produced by the activity of this neuronal network. Our simulation is a *compartmental model*, in which the electrical activity of one or more neurons is simulated by modeling the neuron(s) as a series of small cylindrical segments over which the membrane potential and currents can be taken as approximately constant. The evolution of the membrane potential is given by the cable equation:

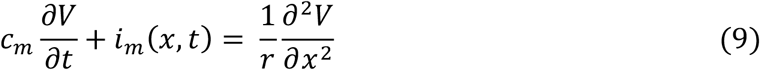

Where *c*_*m*_ is the membrane capacitance per unit length, *r* is the axial resistance of the compartment per unit length, *V* is the membrane potential, and *i*_*m*_ is the current per unit length entering or leaving the compartment through the membrane. Each neuronal compartment is subject to Kirchhoff’s current law:

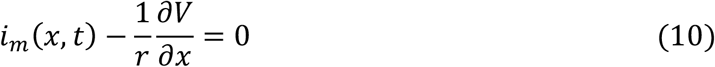

Which states that the net current entering or leaving the compartment must vanish (the second term on the left-hand side represents the net current loss to adjacent segments).

NEURON also has the capability to model ion channels and synapses (which contribute to *i*_*m*_) defined by ordinary differential equations of arbitrary complexity.

We constructed a compartmental model of the neurons in one column of rat sensory cortex based on publicly available data from the Blue Brain Project (BBP)^28^. This data included the spatial location and connectivity matrix of all cells in the column, as well as tuned and experimentally validated models of individual neurons from all cortical laminae including their electrical characteristics (ion channel models and associated parameters such as spatially varying ionic conductances, membrane resistance/capacitance, etc.) and their detailed morphologies based on full reconstructions of neurons observed experimentally (or algorithmically generated clones thereof), reflecting the full diversity of neurons known to be found in rat somatosensory cortex. To save computation time, we instantiate 80% of the cells in the model, selected at random. We confirmed that there was no difference in the spectrum between 80% and 100%. The BBP dataset provides 5 reconstructed (or cloned) morphologies for each of 207 distinct cell types^28^. Each neuron in our model is represented by one of the 5 morphologies for that neuron’s cell type, chosen at random and rotated by a random angle about a line passing through the cell’s soma parallel to the longitudinal axis of the column.

Neurons in our simulated column are innervated by synapses from three populations: 1.) *N*_*thal*_ = 5000 excitatory thalamic neurons conveying feed-forward sensory input (*thalamocortical* connections) modeled as rate-modulated Poisson spike trains, 2.) *N*_*bke,e*_ = 25000 excitatory and *N*_*bke,i*_ = 25000 inhibitory background cortical neurons from other columns (*external cortico-cortical* connections) also modeled as Poisson spike trains, and 3.) other neurons in the simulated column (*internal cortico-cortical* connections). The rate constant of the Poisson processes generating thalamocortical spike trains increases from a baseline rate of 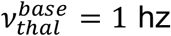 to a stimulus-induced rate of 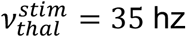 for 50 ms out of every 1000ms (onset and offset are cosine ramps 5 ms in duration), reflecting the temporal structure of tone pips in our experimental preparation, while the rate constant of the external cortico-cortical spike trains remains constant at *v*_*bkg*_ = 7 hz for the duration of the simulation. Thalamic synapses are distributed within the column in a depth-dependent manner, with peaks at 670 μm and 1300 μm below the cortical surface. Synapses from background neurons are formed on neuronal segments in the simulated column with probability proportional to each segment’s surface area.

Synapses from all populations produce membrane currents *i*_*syn*_(*t*) according to

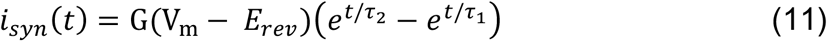

Where *t* is the time since the synapse was activated, *V*_*m*_ is the membrane potential, *E*_*rev*_ is the reversal potential of the synapse, *G* is the weight (max conductance) of the synapse, which is randomly drawn from a lognormal distribution with different center and spread for each input source, and *τ*_1_ and *τ*_2_ are the time constants of the exponential activation/deactivation of the synapse. The values of these parameters for different types of synapses is given in the table below:

| Synapse type | $\tau_1$(ms) | $\tau_2$(ms) | $E_{rev}$ (mV) |
| --- | --- | --- | --- |
| AMPA (e → e) | 1.0 | 3.0 | 0 |
| AMPA (e → i) | 0.1 | 0.5 | 0 |
| GABA (i → e) | 2.7 | 15.0 | -70 |
| GABA (i → i) | 0.2 | 8.0 | -70 |

We applied a modest amount of hand-tuning of these parameters to achieve reasonable baseline firing rate (3-10 Hz) during time periods when the thalamocortical spike trains fire at 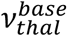 and reproduce the experimentally observed sharp transient stimulus-evoked response (after the transition to 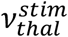) within the simulated column.

### CSEP of the simulated column

The Line Source Approximation (LSA) is used to simulate the extracellular potential at the cortical surface due to the transmembrane currents in each segment of each neuron in the simulation, assuming an isotropic and purely Ohmic extracellular medium^58^. These contributions from individual neuronal segments are summed to compute the total CSEP due to the entire simulated column:

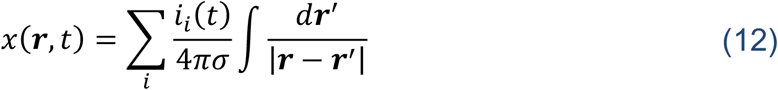

Where *x*(***r***, *t*) is the extracellular potential, *i*_*i*_(*t*) is the current going through neuronal segment *i* at time *t, σ* = 0.3 S/m is the conductivity of the extracellular medium, and the variable of integration ***r***′ runs from one end of segment *i* to the other. The sum in equation (12) runs over all segments *i* in the simulated column. To account for the nonzero spatial extent of the μECoG electrode, we compute *x*(***r***, *t*) at 100 randomly and uniformly sampled points within the 20 μm radius of the μECoG electrode and average them.

### Dependence of response magnitude and frequency on input amplitude

We determined the relationship between z-scored response magnitude (between 10-200 Hz) and the frequency of CSEPs (between 10-200 Hz) by varying the magnitude of inputs in both the experimental data and in the simulations. For simulation data, we varied the frequency 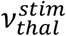 of the feed-forward thalamocortical input and measured the frequency at the maximum of the mean (across stimuli) Z-scored response. Each of the 8 simulations shown in figure 5 is an average over 20 stimuli, performed at 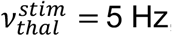, 10 Hz, 15 Hz, 26 Hz, 35 Hz, 44 Hz. For experimental data, at each tuned electrode, we varied the input amplitude (stimulus attenuation) of the best-frequency in the tone stimuli, and measured the frequency at the maximum of the mean (across stimuli) Z-scored response. To compare/combine data, the frequency and responses were normalized to [0 1] for each electrode, and across the simulations. For the experimental data, we only included auditory stimulus attenuations that fell within the FRA to prevent floor effects, and only include tuned channels that had a maximum response above 3 (z-scored) and an FRA boundary that included more than 2 attenuations (as this is the variable that is being ‘manipulated’, i.e., the independent variable). Because of this set of selection criteria, for the experimental data, each of the 6 attenuations had a different number of samples (electrodes) included. Thus, in **Figure 5c**, the grey-to-black points (experimental data) contain, from left-to-right [(attenuation) sample size]: (−50db) N = 206, (−40db) N = 259, (−30db) N = 289, (−20db) N = 299, (−10db) N = 299, (0db) N = 299.

### Laminar lesions and isolations

By summing only neuronal segments belonging to neurons in an individual cortical layer, we obtain the *in silico* contributions *x*_*j*_(***r***, *t*) to the CSEP from distinct layers:

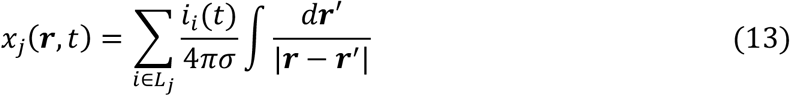

Where *j* ∈ [1,6] denotes the layer whose contribution to *x*(***r***, *t*) is represented by *x*_*j*_(***r***, *t*), and *L*_*j*_ is the set consisting of all neuronal segments comprising neurons in layer *j*. The full signal *x*(***r***, *t*) is the sum of these contributions:

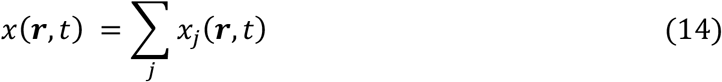

Similarly, we perform the superposition of LSA-computed potentials *excluding* those segments belonging to neurons in a particular layer:

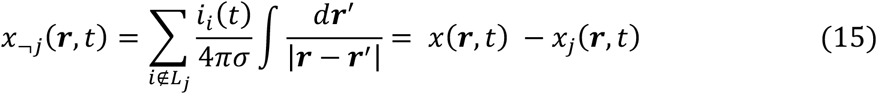

Finally, by summing only the neuronal segments located within a particular range of depths below the surface, we obtain the *in silico* contributions to the CSEP from 200 μm slices of the column:

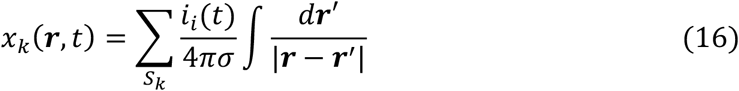

Where *S*_*k*_ := {*i*: *z*_*i*_ ∈ [*k, k* + 1] × 200 μm} is the set of neuronal segments whose midpoints are between 200*k* and 200(*k* + 1) μm below the surface. Note that most neurons’ dendritic arbors extend beyond the slice boundaries, therefore each slice contains segments from neurons in a multitude of layers, and a given neuron may contribute to multiple slices.

### Normalization of simulated CSEP components to baseline

To assess the contributions of sources at different depths to CSEPs (Fig.6,7), we would like to normalize the CSEP contributions to baseline in a way that preserves their relative magnitudes. Using the z-score of each contribution to its own baseline does not preserve relative magnitudes between contributions. For example, two contributions that differ only by a constant multiplicative factor will have the same z-score. That is, if *x*_*j*_(*t*) = *c* · *x*_*i*_(*t*) for two layers *i* and *j*, and for some constant *c*, then the z-scores of these two contributions are

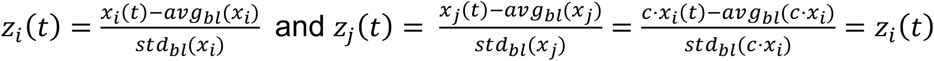

(where *avg*_*bl*_(*x*) and *std*_*bl*_(*x*) denote the average and standard deviation, respectively, of *x* during the baseline periods – the silence between tone pips) whereas we would like a normalization procedure that gives *z*_*j*_(*t*) = *c* · *z*_*i*_(*t*) in such a case. To achieve this, in Figure 6 & 7, we use the *ratio* of each contribution during the stimulus to the *total* simulated baseline signal. The ratio *r*_*i*_ for a contribution *x*_*i*_(*t*) (representing an anatomical layer or 200 μm slice) is then given by

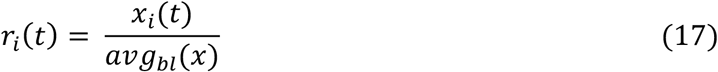

As with the z-score analysis used for the full CSEP (both in experiments and simulations, Fig.4,5), this normalization is done independently for each neural frequency bin. In Supplementary Figures 2 & 3, we show that our conclusions are robust to the choice of normalization.

### Dependence of simulated CSEP contributions on number of segments/neurons, depth, and synchronicity

The CSEP contribution from a given subset of the column (anatomically defined layer, or 200 μm slice) will depend on the number of neuronal segments in the subset, the distance of those neurons from the recording electrodes, and the synchronicity of those neurons’ activity. To determine the relative importance of these three factors, we performed an L_2_-regularized regression to fit the magnitude of each contribution’s high-gamma peak (the maximum of each normalized CSEP contribution across frequency bins) as a linear function of the number of simulated neurons in each layer, or the number of neuronal segments in each 200 μm slice, the average depth of the contributing segments below the surface, and the synchronicity between somatic membrane potentials, defined as the average of Pearson’s correlation coefficient of the membrane potentials over all pairs of somas in the subset. Each of the 3 independent variables, and the dependent variable (high-gamma contribution magnitude) was normalized by dividing by the maximum across layers or slices. The L_2_ regularization parameter was chosen to be *α* = 0.01. The magnitude of the fit coefficients gave the relative importance of each factor in determining the magnitude of the CSEP contributions in our model.

## Data and Software Availability

The datasets and software are available upon request.

## Supplemental Information for

**SFig. 1.**
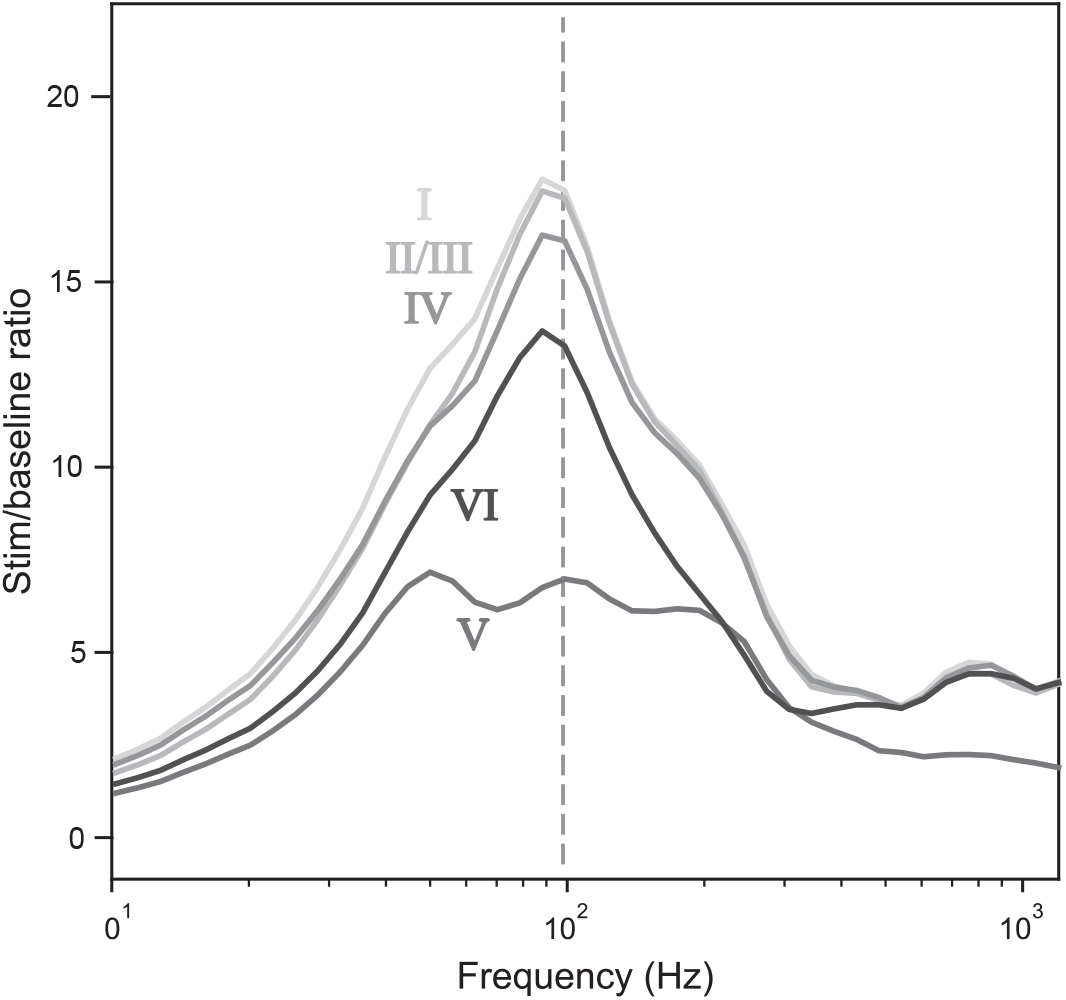
High gamma peak is greatly diminished when layer V is lesioned. We computed the simulated CSEP from the column with each layer lesioned (Methods, main text). Each trace in **SFig.1** shows the difference between the full CSEP and the corresponding trace in Figure 6b in the main text. The high gamma peak survives lesioning of all layers except layer V, suggesting that this component of the signal originates primarily in layer V neurons. Lesion of layer V produced the sharpest decrease in CSEP amplitude across almost the entire range of frequencies analyzed.

**SFig. 2.**
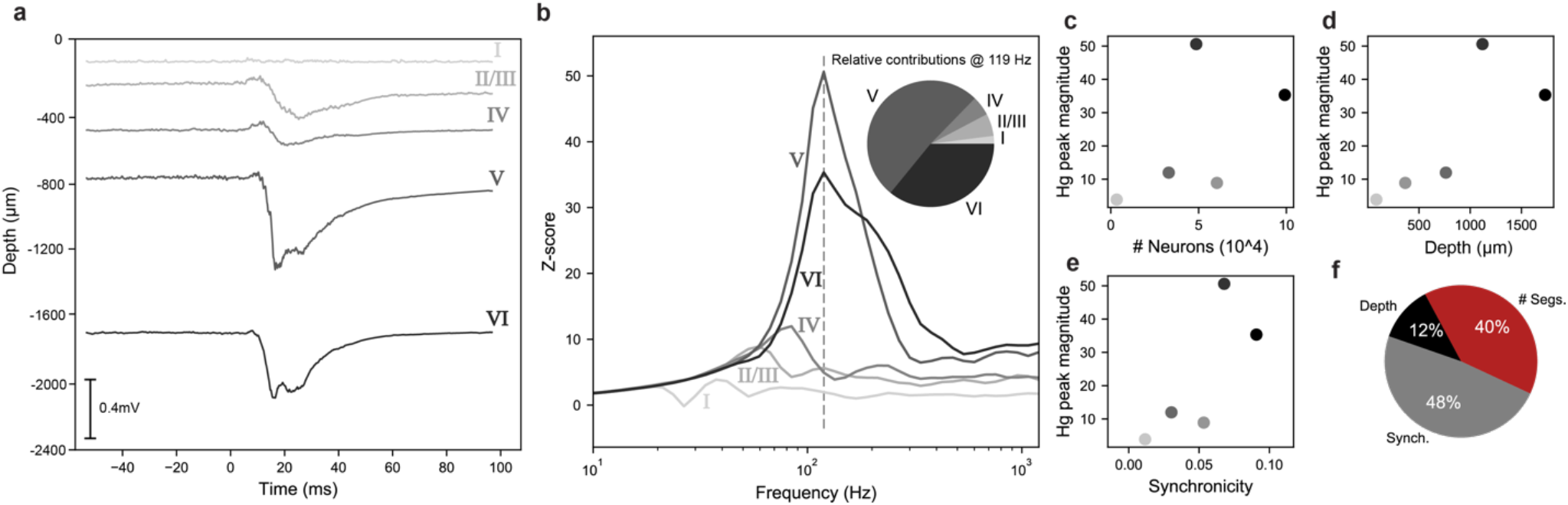
Z-scored evoked responses are strongest in infragranular layers. Due to difficulties interpreting z-scored evoked laminar contributions relative to baseline, in **Figures 6**-**7** we applied a ratio-based normalization procedure in which the stimulus-evoked activity is simply divided by the baseline average activity level (see Methods). While this allows us to accurately capture the relative magnitudes of each of the laminar contributions, information about which layers respond strongest *above their baseline level* remains hidden when the normalization factor (denominator in equation (19)) is not layer-specific. To address this, we revert to the z-score normalization procedure, z-scoring each laminar contribution to that layer’s baseline statistics (mean and standard deviation during interstimulus silence), then reproduce **Figure 6**. The z-scored evoked responses are shown in **SFigure 2b**, where layers V and VI can be seen dominating the high gamma peak, with layers I – IV contributing minimally, as observed in **Figure 6b**. This suggests that in addition to contributing the largest fraction of the CSEP, layers V and VI also have by far the largest evoked response above baseline. Still, most layers, excepting layer I, exhibit strong responses several standard deviations above baseline levels. Thus, although all layers are stimulus-driven to an extent, layer I is the least stimulus-driven, which may owe to its relative lack of input (**Figure 4b**). Finally, the frequency of the peak in the infragranular layers is slightly higher than when using ratio normalization.

**SFig. 3.**
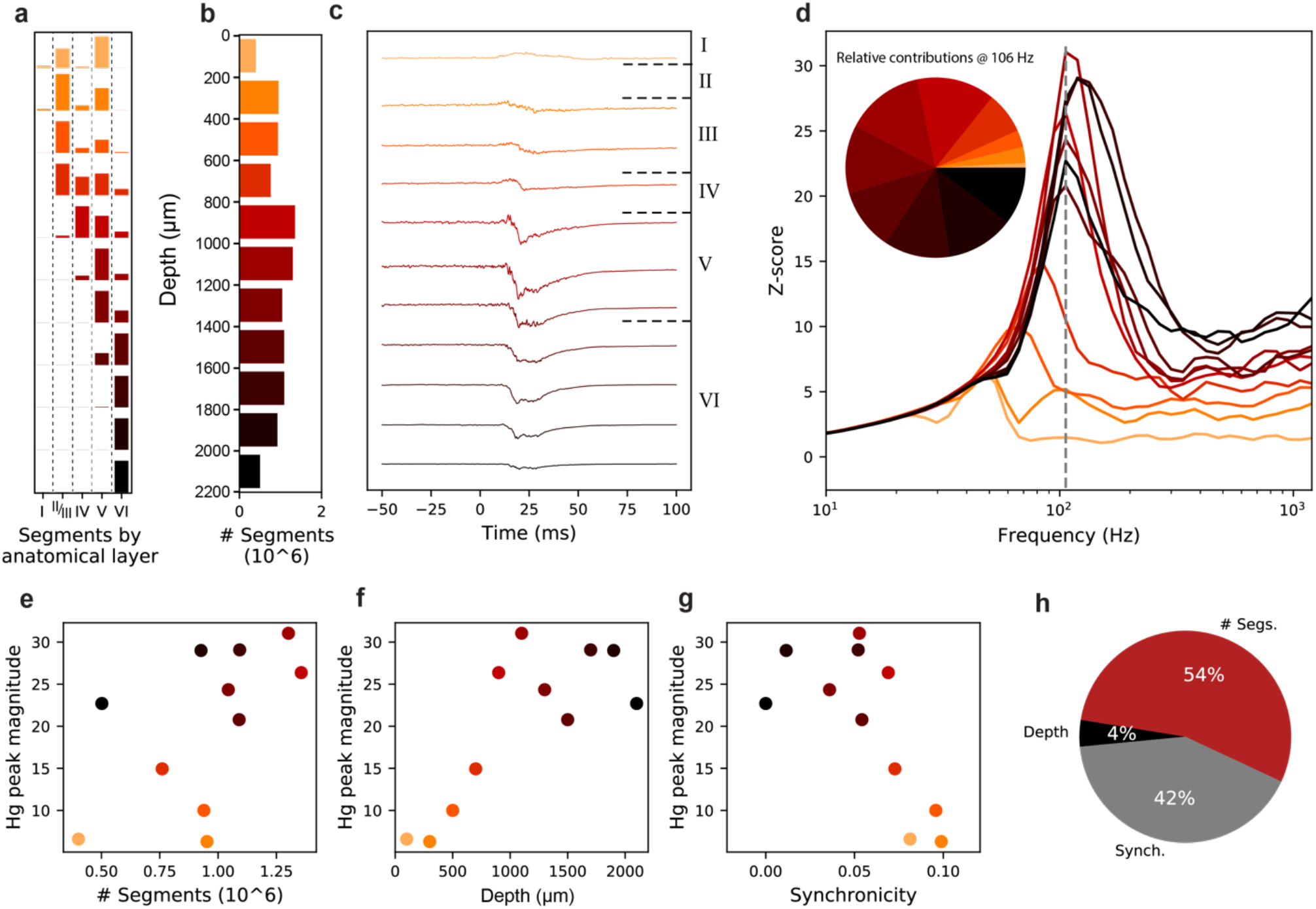
Z-scored evoked responses are strongest in infragranular layers. In **SFigure 1**, we analyzed each layer’s contribution to the stimulus evoked CSEP relative to that layer’s activity during baseline. Here, we apply the same technique to the CSEP contributions from 200μm slices of the column, normalizing contributions to their own baseline statistics instead of using equation (19), thereby sacrificing information about the relative magnitudes of the slices’ contributions in favor of information about magnitude of each slice’s response above baseline. In **SFigure 3d**, the z-scored evoked responses of the slices are shown. As seen in **SFigure 1**, the peak of the response in most slices is found at a slightly higher frequency. Contributions to the high gamma peak at 106 Hz are more equally divided across the slices when looking at this z-scored data as compared to the ratio-normalized data shown in **Figure 7d**. However, infragranular layers are still seen to respond more strongly above baseline than layers I – IV, and the biophysical parameters are of similar proportions (**SFig.3e**).

## Notes

### Competing Interest Statement

The authors have declared no competing interest.

